# Engineering acidophilic non-conventional yeasts as low-pH cell factories for organic acid production

**DOI:** 10.64898/2026.11.24.768970

**Authors:** Reem Swidah, Federico Visinoni, Soukaina Timouma, Angus Nelson, Maira Hernandez-Guzman, Fernando Valle, Daniela Delneri

## Abstract

Acidophilic non-conventional yeasts (NCYs) thrive at low pH and offer promising hosts for organic acid production, but their exploitation is constrained by the limited availability of genetic engineering tools. Here, we optimised a chemical transformation protocol enabling efficient DNA delivery into several NCYs, including *M. bulderi*, *S. bombicola* and *S. batistae, K. aerobia*, *K. naganishii* and *Y. lipolytica*. We also developed a Pan/ARS-containing expression vectors compatible with YeastFab for one-pot assembly of standardised promoters, open reading frames (ORFs) and terminators. As proof of concept, we engineered *M. bulderi*, *S. bombicola* and *S. batistae* for lactic acid production. Codon-optimised *ldh* genes from *B. megaterium* and *L. helveticus* were combined with regulatory elements (REs) from *S. cerevisiae* or *M. bulderi* and inserted episomally in the host. Lactic acid production varied substantially among hosts and constructs, with the highest amount achieved in *S. batistae* (ca 6.4 g/L lactic acid) expressing *B. megaterium ldh* under the control of *S. cerevisiae* Res. To enable targeted genome engineering, we developed a CREEPY-derived CRISPR–Cas9 system supporting multiplex gRNA assembly, achieving over 94% gene deletion efficiency in *S. bombicola*. In *Starmerella* species, Aro10p contributes to the conversion of pyruvate to acetaldehyde, potentially diverting carbon away from lactic acid production. Deletion of *ARO10 in S. bombicola* increased lactic acid production from both endogenous and exogenous *ldh* pathways up to 6 g/L. Notably, S. *bombicola* was unable to grow on lactate as the sole carbon source, whereas *S. batistae* utilised both D- and L-lactate, highlighting *S. bombicola* as a potentially advantageous production host by limiting product re-assimilation. Together, these genetic tools and metabolic engineering strategies expand the engineering capabilities of acidophilic NCYs and establish a foundation for their development as low-pH cell factories for the sustainable biomanufacture of organic acids and other bio-based chemicals.

**Graphical abstract Figure:** An integrated synthetic biology pipeline for engineering acidophilic non-conventional yeasts for sustainable lactic acid production.
Acidophilic non-conventional yeast (NCY) hosts were first identified for their ability to grow under low-pH conditions, and an efficient chemical transformation platform was subsequently developed to enable plasmid delivery. A versatile synthetic biology toolkit, comprising broad-host-range Pan/ARS expression vectors and the CRISPR/Cas9 CREEPY genome-editing system, was established to facilitate genetic engineering. YeastFab technology was subsequently used to construct heterologous *ldh* expression cassettes, enabling lactic acid biosynthesis and revealing species-specific responses to *ldh* expression. Finally, metabolic engineering via *ARO10* deletion further enhanced lactic acid production in *S. bombicola*, establishing next-generation acidophilic yeast cell factories for sustainable biomanufacturing.

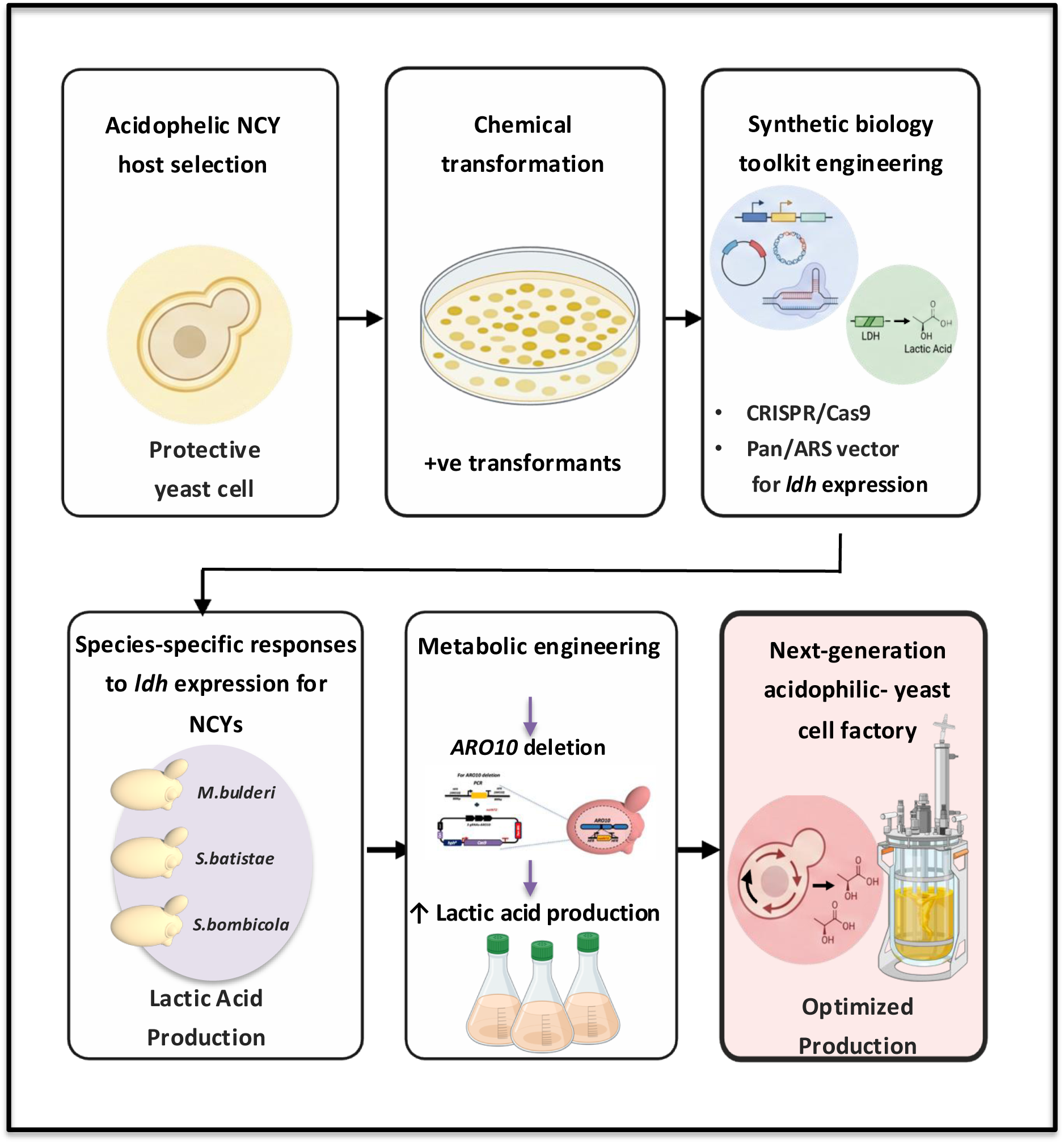

## Introduction

Organic acids are key bio-based platform chemicals used in the food, pharmaceutical, cosmetic, and chemical industries. Lactic acid is especially important because of the rapid growth of the poly (lactic acid) (PLA) market, as PLA is one of the most commercially successful biodegradable polymers. Demand for sustainable materials and concern over fossil-derived plastics have driven interest in economically viable microbial fermentation platforms for lactic acid (Atasoy, Álvarez Ordóñez et al. 2024, Wang, Cui et al. 2025). Future progress will depend not only on strain performance but also on more efficient fermentation and downstream processing (Tsaruk, Filip et al. 2025, Wang, Cui et al. 2025).

Yeasts are attractive organic acid hosts because they are robust under industrial conditions, tolerate osmotic stress, use diverse carbon sources, and have advanced genetic tools available (Pedelacq, Cabantous et al. 2006, Tran and Zhao 2022, Tsaruk, Filip et al. 2025, Yook and Alper 2025). Systems and synthetic biology, CRISPR-based genome editing, and adaptive laboratory evolution (ALE) have expanded the engineering of conventional and non-conventional yeasts, positioning them as next-generation cell factories for organic acids and other value-added biochemicals (Liu, Jin et al. 2023, Yook and Alper 2025).

*Saccharomyces cerevisiae* remains the most extensively engineered yeast for lactic acid production. Strategies include heterologous lactate dehydrogenase expression, elimination of competing pyruvate-consuming pathways, transporter engineering, and redox balancing (Zhu, Luo et al. 2022, Altvater, Tomico-Cuenca et al. 2025). Combining ALE with rational engineering has yielded strains producing over 100 g L⁻¹ lactic acid under industrially relevant conditions (Zhu, Luo et al. 2022, Altvater, Tomico-Cuenca et al. 2025), demonstrating the importance of *S. cerevisiae* for large-scale production.

Lactic acid toxicity remains a major physiological limitation. Undissociated acid diffuses through the plasma membrane and dissociates intracellularly, acidifying the cytoplasm; maintaining pH homeostasis then costs substantial ATP, reducing growth and production. Improving tolerance to low pH and organic acid stress is therefore a central goal in industrial strain development, and critical determinants of success (Pangestu, Kahar et al. 2022, Tran and Zhao 2022).

Controlling fermentation at neutral pH around 7.0 is a major industrial challenge because it requires the continuous addition of neutralising agents, which reduces acid inhibition but generates lactate salts that must be converted to free lactic acid downstream, increasing operating costs, raw material use, and waste, so processes operating under acidic conditions can substantially improve economic viability (Wang, Cui et al. 2026).

Low-pH fermentation is therefore a key objective for next-generation lactic acid bioprocesses. It reduces or eliminates neutralising agents, minimises salt formation and wastewater, and simplifies recovery of the free acid (Atasoy, Álvarez Ordóñez et al. 2024, Wang, Cui et al. 2026). It also lowers contamination risk, improving robustness and reliability. Acid-tolerant microorganisms are considered a key strategy for making future biorefineries more sustainable and competitive (Atasoy, Álvarez Ordóñez et al. 2024, Wang, Cui et al. 2026).

Non-conventional yeasts further expand opportunities for low-pH organic acid production. Species such as *Pichia kudriavzevii* (*Issatchenkia orientalis*), *Yarrowia lipolytica*, *Kluyveromyces marxianus*, and other acid-tolerant yeasts naturally resist acidic environments and fermentation inhibitors while maintaining efficient carbon metabolism (Fickers, Cheng et al. 2020, Wang, Cui et al. 2025, Yook and Alper 2025). Non-conventional yeasts are regarded as promising platforms for low-cost, sustainable organic acid production under conditions that inhibit many traditional hosts (Wang, Cui et al. 2025, Yook and Alper 2025).

Over the past decade, experimental findings have rapidly strengthened this view. Acid-tolerant *P. kudriavzevii* strains have reached high lactic acid titers at low pH (Park, Bae et al. 2018), and engineered strains of this species have also efficiently converted mixed sugars to lactic acid with high productivity at low pH (Lin, Kim et al. 2026) [17]. Most of the lactic acid produced commercially by fermentation nowadays is based on the use of LAB, *S. cerevisiae* and *Pichia kudriavzevii* strains.

Among the emerging non-conventional yeasts, *Maudiozyma bulderi* (formerly *Kazachstania bulderi*), *Starmerella bombicola*, and *Starmerella batistae* have attracted increasing interest because of their high tolerance to acidic environments and their potential for sustainable industrial biotechnology (Balarezo-Cisneros, Timouma et al. 2023, Wang, Wang et al. 2025). *M. bulderi* is particularly attractive as it can grow efficiently over a broad pH range, including highly acidic conditions (pH 2.5–3.0), making it a promising host for low-pH organic acid fermentations. Similarly, *Starmerella* species possess unique metabolic capabilities, including the natural production of industrially important metabolites and an inherent ability to thrive under acidic conditions, making them attractive candidates for next-generation microbial cell factories (Wang, Wang et al. 2025, Timouma, Hanak et al. 2026). Recent advances in long-read genome sequencing and genome annotation have provided valuable genomic resources for these species, creating new opportunities for strain engineering and metabolic optimisation (Balarezo-Cisneros, Timouma et al. 2023; Timouma, Hanak et al. 2026). Despite these advances, efficient genetic engineering of these yeasts remains challenging because of the limited availability of transformation methods and molecular tools.

Despite their considerable industrial potential and the increasing availability of high-quality genome sequences, most non-conventional yeasts remain genetically intractable. Efficient transformation methods, expression vectors, regulatory elements, selectable markers and robust genome editing systems remain unavailable or poorly developed for many emerging yeast species, limiting both biological studies and strain engineering. In particular, low transformation efficiency and the lack of reliable synthetic biology toolkits continue to hinder the engineering of promising acid-tolerant yeasts, including *M. bulderi*, *S. bombicola*, and *S. batistae*, restricting their wider adoption as industrial production hosts (Moon, An et al. 2025, Yook and Alper 2025). These limitations highlight the need for robust molecular tools that enable rapid and efficient engineering of these emerging yeast cell factories.

Here, we addressed these challenges by optimising transformation protocols and developing a comprehensive synthetic biology toolkit for *M. bulderi*, *S. bombicola*, and *S. batistae*. We established an efficient transformation platform, identified suitable selectable markers, and developed a broad-host-range expression vector, that enables stable plasmid maintenance and heterologous gene expression across these acid-tolerant yeast species. In addition, we engineered the CRISPR/Cas9-based CREEPY genome editing system, originally developed for *Saccharomyces cerevisiae*, to enable its stable maintenance in non-conventional yeasts (Zhao, Coelho et al. 2023), and demonstrated multiplexed guide RNA expression to improve genome editing efficiency and accelerate strain engineering. Together, these tools expand the genetic accessibility of these previously intractable yeasts and provide a versatile platform for developing next-generation microbial cell factories for the sustainable production of lactic acid and other organic acids.

## Materials and methods

### Strains, plasmids, synthetic DNA fragments, and growth media

The yeast strains used in this study included *Saccharomyces cerevisiae* BY4741, *Maudiozyma bulderi* CBS 8639, *Starmerella batistae*, and *Starmerella bombicola*, all obtained from the laboratory of Daniela Delneri (Manchester Institute of Biotechnology, The University of Manchester, UK). YeastFab plasmids (pHCKan-P, pHCKan-O, pHCKan-T, and pPOT-LEU2) were obtained from the laboratory of Yizhi Cai (Manchester Institute of Biotechnology, The University of Manchester, UK; Guo et al., 2015). The broad-host-range expression vector backbone was derived from pOT2-*URA3* (Guo et al., 2015), and the pan/ARS replication origin was derived from pUDBzB-41 (Jayaprakash, Barroso et al., 2023). The kanMX (G418 resistance) cassette was obtained from the laboratory of Mark Ashe (Faculty of Biology, Medicine and Health, The University of Manchester, UK; Swidah, Wang et al. 2015). The CRISPR/Cas9 expression creepy system derived from Jef D. Boeke — Institute for Systems Genetics, NYU Grossman School of Medicine, New York University, New York, NY, USA (Zhao, Coelho et al. 2023). The L-lactate dehydrogenase (*ldh*) gene from *Lactobacillus helveticus* and the D-lactate dehydrogenase (*ldh*) gene from *Bacillus megaterium* were codon-optimised for yeast expression and synthesised by Twist Bioscience (USA) in standard cloning vectors carrying a kanamycin resistance marker. Yeast strains were routinely cultured in yeast extract-peptone-dextrose (YPD) medium (10 g L⁻¹ yeast extract, 20 g L⁻¹ peptone, and 20 g L⁻¹ glucose). Selection was performed on synthetic complete dextrose (SCD) medium supplemented with 2% glucose and the appropriate selective agent, including G418 (Geneticin; Thermo Fisher Scientific) or amino acid dropout medium, where appropriate. *Escherichia coli* strains were cultured in Luria–Bertani (LB) medium and selected with carbenicillin (100 μg mL⁻¹), kanamycin (50 μg mL⁻¹), as appropriate. Unless otherwise stated, all media components were purchased from Formedium (UK) or Fisher Scientific (UK).

### Bacterial transformation

Plasmids were transformed into chemically competent *Escherichia coli* DH5α cells by heat shock following standard procedures. For plasmids containing the ccdB counter-selection cassette, chemically competent ccdB-resistant *E. coli* cells were used instead. Briefly, competent cells were incubated with plasmid DNA on ice for 30 min, heat shocked at 42°C for 45 s, and recovered in pre-warmed S.O.C. medium for 1 h at 37°C with shaking (225 rpm). Cells were concentrated by centrifugation and plated onto LB agar supplemented with the appropriate selective antibiotic, followed by overnight incubation at 37°C. Transformation efficiency was monitored using pUC19 as a positive control. For plasmids smaller than 10 kb, DH5α Mix & Go! Competent Cells (Zymo Research, Orange, CA, USA) were used according to the manufacturer’s instructions, except that the competent cells were prepared at approximately threefold higher concentration than recommended to improve transformation efficiency. When transforming plasmids containing the ccdB cassette, ccdB Survival Competent Cells (Zymo Research) were used according to the manufacturer’s instructions.

### Chemical transformation of non-conventional yeasts

Chemical transformation of non-conventional yeasts was performed using an optimised lithium acetate/polyethylene glycol (LiAc/PEG) protocol. Overnight yeast cultures were grown in yeast extract-peptone-dextrose (YPD) medium at 30°C with shaking (250 rpm). Cultures were diluted to an initial optical density at 600 nm (OD₆₀₀) of 0.2 in 25 mL fresh YPD medium and grown to an OD₆₀₀ of approximately 1.0. Cells were harvested by centrifugation at 4,000 rpm for 5 min at room temperature, washed twice with sterile water, and resuspended in 1 mL of Solution I (50 mM lithium acetate, 10 mM Tris-HCl, 1 mM EDTA, pH 7.5). For each transformation, 100 μL of competent cells were mixed with 100 μg of denatured salmon sperm DNA and 1 μg of plasmid or linear DNA. 100 μg salmon sperm DNA was denatured at 95°C for 5 min and immediately chilled on ice before use. Negative control transformations lacking transforming DNA were included in all experiments. Subsequently, 600 μL of Solution II (40% PEG 4000, 50 mM lithium acetate, 10 mM Tris-HCl, 1 mM EDTA, pH 7.5) was added, and the mixtures were vortexed vigorously for 1 min before incubation at 30°C with shaking (250 rpm) for 90–120 min. Following transformation, cells were collected by centrifugation at 13,000 rpm for 5 min, resuspended and recovered in 1ml liquid YPD medium for 24 h at room temperature, an optimisation that significantly improved the recovery of transformed non-conventional yeasts. Cells were then harvested by centrifugation, resuspended in 200 μL of sterile water, and spread onto the appropriate selective agar medium. Plates were incubated at 30°C for 3–4 days until transformant colonies became visible. Solution I: 50 mM lithium acetate, 10 mM Tris-HCl, 1 mM EDTA (pH 7.5). Solution II: 40% (w/v) PEG 4000, 50 mM lithium acetate, 10 mM Tris-HCl, 1 mM EDTA (pH 7.5). This protocol was adapted from the lithium acetate transformation method described by Lodens et al (Lodens, De Graeve et al. 2018) and further optimised for the efficient transformation of acid-tolerant non-conventional yeasts.

### Construction of a YeastFab-compatible broad-host-range expression vector

A compact (∼6 kb) YeastFab-compatible broad-host-range destination vector was engineered for use throughout this study. The vector was designed to support plasmid propagation in *Escherichia coli*, replication in both conventional and non-conventional yeasts, and modular assembly of synthetic genetic cargo for interkingdom DNA transfer. The vector backbone comprised a pan-fungal autonomously replicating sequence (Pan/ARS) to enable plasmid maintenance in *Saccharomyces cerevisiae* and diverse non-conventional yeast species (Jayaprakash, Barroso et al. 2023), a *kanMX* cassette conferring G418 resistance for yeast selection (Swidah, Wang et al. 2015), and an AmpR/ColE1 module for plasmid propagation and selection in *E. coli*. To facilitate modular cloning, a *ccdB* counter-selection cassette flanked by Type IIS restriction enzyme recognition sites was incorporated, enabling replacement with standardised YeastFab promoter, open reading frame (ORF), and terminator modules using Golden Gate assembly (Guo, Dong et al. 2015, Garcia-Ruiz, Auxillos et al. 2018). This modular design enables rapid construction of both single transcriptional units and multigene pathways using standardised YeastFab DNA parts. Rather than using conventional restriction-ligation cloning, the vector backbone was assembled directly in vivo in *S. cerevisiae* through homologous recombination. Individual DNA fragments were generated with approximately 60 bp homologous overlaps at both termini to direct recombination with adjacent fragments, enabling seamless assembly of the complete plasmid. Successful assembly was confirmed by three junction PCR assays spanning adjacent assembly junctions. Five independent transformants were analysed, and all produced the expected amplicons, demonstrating efficient and reproducible plasmid assembly. Following assembly, plasmids were recovered from yeast using the previously described EASY-C (Extraction and Analysis of Small Yeast Chromosomes) workflow (Swidah, Monti et al. 2026). Briefly, plasmid DNA was extracted from yeast, transformed into *ccdB*-resistant *E. coli*, propagated under ampicillin selection, purified using a commercial plasmid purification kit, and verified by Sanger sequencing. Only sequence-verified plasmids were used for all subsequent experiments. The sequence-verified plasmid served as a reusable YeastFab-compatible broad-host-range destination vector for the insertion of standardised genetic modules and biosynthetic pathways.

### Construction of a broad-host-range CRISPR/Cas9 CREEPY shuttle vector

A broad-host-range CRISPR/Cas9 shuttle vector compatible with the CREEPY genome editing system was constructed by in vivo assembly in *Saccharomyces cerevisiae* through homologous recombination. The vector was assembled from four PCR-amplified DNA fragments. The first fragment comprised the Pan/ARS, a *TEF1* promoter-driven *Cas9* expression cassette, and the *PHO5* terminator, amplified from pUDPzb_U1 (Jayaprakash, Barroso et al. 2023). The second fragment contained the bacterial origin of replication (ColE1 ori) and the *AmpR* cassette for plasmid propagation and selection in *Escherichia coli* (Jayaprakash, Barroso et al. 2023). The third fragment comprised the *natNT2* nourseothricin resistance cassette driven by the *TEF1* promoter and terminated by the *ADH1* terminator, amplified from pZC2 (Carter and Delneri 2010). The fourth fragment contained a GFP reporter flanked by BsmBI recognition sites, enabling Golden Gate assembly of single or multiplex guide RNA (gRNA) expression cassettes. Insertion of the gRNA expression cassette replaced the GFP reporter, allowing rapid identification of correctly assembled constructs through loss of green fluorescence (Pedelacq, Cabantous et al. 2006, Zhao, Coelho et al. 2023). The four DNA fragments were PCR-amplified, purified, mixed in equimolar amounts, and co-transformed into *S. cerevisiae* for in vivo assembly by homologous recombination. Each fragment contained approximately 60 bp homologous overlaps with adjacent fragments to facilitate seamless plasmid assembly. Transformant harbouring the right construct was selected in *natNT2 100* µg/ml. Correct assembly was confirmed by three junction PCR assays spanning adjacent assembly junctions. Plasmids were subsequently recovered from yeast using the previously described EASY-C (Extraction and Analysis of Small Yeast Chromosomes) workflow (Swidah, Monti et al. 2026), transformed into *E. coli* DH5α, and verified by full-plasmid sequencing (Plasmidsaurus, USA). To ensure compatibility with Golden Gate cloning, the internal BsmBI recognition site within the Cas9 coding sequence was removed by synonymous site-directed mutagenesis using the Q5® Site-Directed Mutagenesis Kit (New England Biolabs, Ipswich, MA, USA) according to the manufacturer’s instructions, without altering the encoded amino acid sequence. The mutagenized plasmid was transformed into *Escherichia coli* DH5α and verified by full-plasmid sequencing (Plasmidsaurus, USA). The resulting sequence-verified plasmid functions as a broad-host-range CRISPR/Cas9 shuttle vector that can be propagated in both *E. coli* and non-conventional yeasts, enabling efficient Golden Gate assembly of single or multiplex guide RNA (gRNA) expression cassettes for CRISPR/Cas9-mediated genome editing.

### Recovery of the expression vector using EASY-C

Correctly assembled plasmids were recovered from yeast using the Extraction and Analysis of Small Yeast Chromosomes (EASY-C) workflow (Swidah, Monti et al. 2026). Briefly, plasmid DNA was isolated from yeast, transformed into chemically competent *E. coli* DH5α cells for propagation, and purified using the QIAprep Spin Miniprep Kit (QIAGEN) according to the manufacturer’s instructions.

### Sequence verification

Purified plasmids were verified by full-plasmid Sanger sequencing using the Plasmidsaurus sequencing service. Sequencing reads were aligned to the reference plasmid sequence to confirm complete and accurate plasmid assembly. Only sequence-verified constructs were used for subsequent experiments.

### Generation and cloning of guide RNA (gRNA) expression cassettes

Guide RNA (gRNA) expression cassettes, including the gRNA scaffold sequence, were PCR-amplified from the TOPO vector developed for the CREEPY CRISPR/Cas9 system (Zhao, Coelho et al. 2023). Purified PCR products were assembled into the CRISPR/Cas9 shuttle vector by Golden Gate assembly using BsmBI and T4 DNA ligase (New England Biolabs, Ipswich, MA, USA) according to the manufacturer’s instructions. Recombinant plasmids were transformed into *Escherichia coli* DH5α, and correctly assembled constructs were identified by loss of GFP fluorescence and confirmed by colony PCR and Sanger sequencing.

### YeastFab assembly of transcriptional units

Standardised YeastFab biological parts, including promoter, open reading frame (ORF), and terminator modules, were assembled into the pOT destination vector using a one-pot Golden Gate reaction. Individual DNA parts and the destination vector were combined in equimolar amounts and assembled using BsmBI and T4 DNA ligase (New England Biolabs, Ipswich, MA, USA) according to the manufacturer’s instructions and the YeastFab assembly workflow (Guo, Dong et al. 2015). During assembly, the *ccdB* counter-selection cassette within the destination vector was replaced by the desired transcriptional unit. As expression of *ccdB* is toxic to standard *Escherichia coli* cloning strains, only plasmids in which the *ccdB* cassette had been successfully replaced yielded viable *E. coli* DH5α transformants. Plasmid DNA was extracted from viable colonies using a standard plasmid purification kit and screened by BsaI restriction digestion to confirm the expected plasmid architecture. Correctly assembled constructs were subsequently verified by full-plasmid long-read sequencing (Plasmidsaurus, Eugene, OR, USA), and only sequence-verified plasmids were used for subsequent experiments.

### Colony PCR confirmation

Colony PCR was performed to confirm the correct assembly of plasmids and genetic constructs using GoTaq® Green Master Mix (Promega, Madison, WI, USA) according to the manufacturer’s instructions. Briefly, yeast colonies were lysed in 20 mM NaOH using the sodium hydroxide (NaOH) colony PCR method, and the resulting lysates were used directly as PCR templates. PCR products were separated by agarose gel electrophoresis, stained with SafeStain (NBS Biologicals, Huntingdon, UK), and visualised using a GelDoc™ Go Imaging System (Bio-Rad Laboratories, Hercules, CA, USA).

### Spot assay

Overnight yeast cultures were grown in the appropriate selective medium, adjusted to an OD₆₀₀ of **0.1**, and serially diluted (10-fold). Aliquots (10 μL) of each dilution were spotted onto the appropriate minimal agar medium and incubated at 30°C for 3–4 days before imaging and phenotypic assessment.

### Lactic acid quantification by HPLC

For lactic acid production assays, overnight cultures of each yeast strain were grown in YPD medium supplemented with G418 at the appropriate concentration for each species (200 μg mL⁻¹ for *Maudiozyma bulderi*, 600 μg mL⁻¹ for *Starmerella bombicola*, and 2,000 μg mL⁻¹ for *Starmerella batistae*). Cultures were diluted to an initial optical density at 600 nm (OD₆₀₀) of 0.1 in 30 mL of fresh YPD medium containing the corresponding concentration of G418 and incubated under the appropriate growth conditions. Three independent biological replicates were performed for each strain. Culture samples were collected at the indicated time points, clarified by centrifugation, and the supernatants were filtered through 0.22 μm syringe filters before HPLC analysis. Lactic acid concentrations were determined using a 1260 Infinity II LC System equipped with a refractive index detector (Agilent Technologies, Santa Clara, CA, USA). Separation was performed on a Hi-Plex H column (300 × 7.8 mm; Agilent Technologies) maintained at 55°C using 5 mM H₂SO₄ as the mobile phase at a flow rate of 0.8 mL min⁻¹. Lactic acid was identified by its retention time and quantified using calibration curves generated from authentic standards, as previously described (Visinoni, Zhang et al. 2022)..

### Statistical analysis

All quantitative experiments were performed using three independent biological replicates unless otherwise stated. Data are presented as the mean ± standard deviation (SD). Data organisation and descriptive statistics were performed using Microsoft Excel. For experiments involving multiple engineered strains, differences among groups were assessed by one-way analysis of variance (ANOVA), followed by Dunnett’s multiple-comparisons test to compare each engineered strain with the corresponding empty-vector control. A *P* value < 0.05 was considered statistically significant. Significance is indicated as ns, *P* ≥ 0.05; *, *P* < 0.05; **, *P* < 0.01; ***, *P* < 0.001; and ****, *P* < 0.0001.

## Results

**Acidophilic non-conventional yeasts exhibit exceptional growth under low-pH conditions and represent promising hosts for sustainable lactic acid production.**

To assess acid tolerance across the selected non-conventional yeasts (NCYs), *Maudiozyma bulderi*, *Starmerella bombicola*, and *Starmerella batistae* were evaluated by spot assays over a pH range of 7.0–1.5 at 30°C. Cultures were normalised to an OD₆₀₀ of 0.1 and subjected to 10-fold serial dilutions before spotting onto minimal medium. *Saccharomyces cerevisiae* BY4741 and BY4743 were included as haploid and diploid reference strains, respectively (Fig. 1).

**Figure 1.**
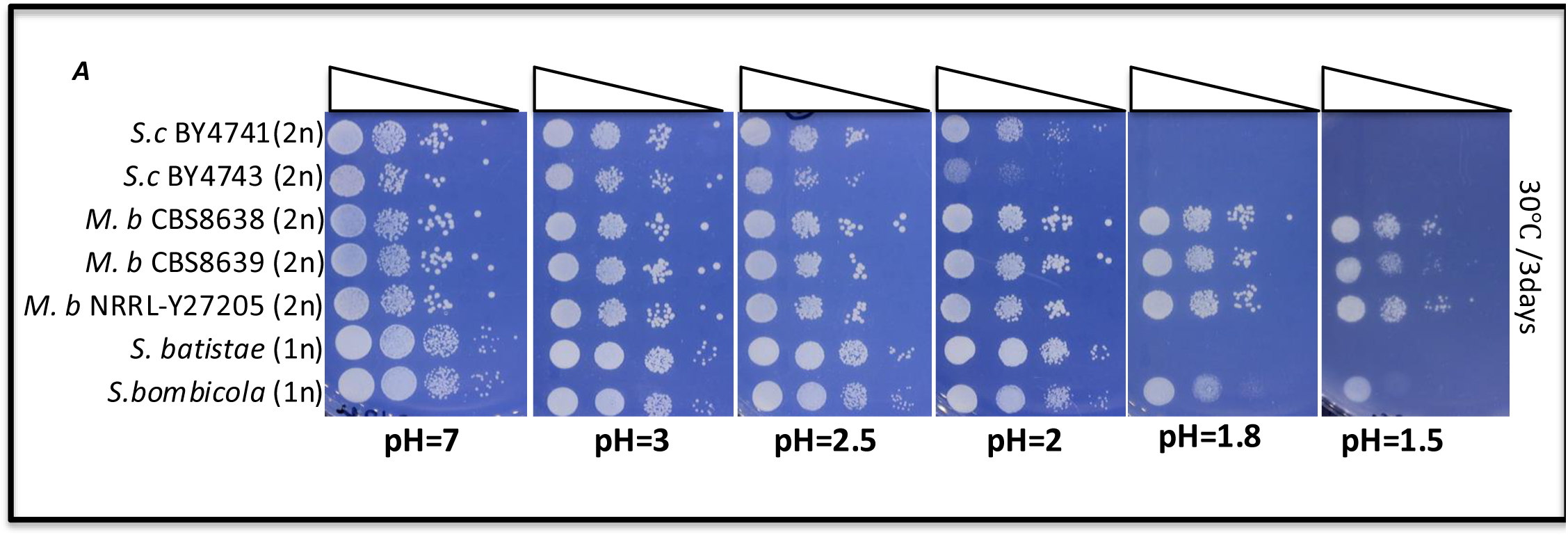
Acid-tolerant non-conventional yeasts exhibit exceptional growth under low-pH conditions and represent promising microbial cell factories for sustainable lactic acid production Representative spot assays showing the growth of *Maudiozyma bulderi*, *Starmerella bombicola*, and *Starmerella batistae* on synthetic complete dextrose (SCD) agar medium at the indicated pH values. Overnight cultures were adjusted to an OD₆₀₀ of 0.1, subjected to 10-fold serial dilutions, and spotted onto SCD agar plates. Plates were incubated at 30°C (A) or 25°C (B) for 3–4 days before imaging.

The selected NCYs exhibited pronounced acid tolerance and, in several cases, continued to thrive under strongly acidic conditions that substantially impaired the *S. cerevisiae* reference strains. Among the strains tested, diploid *M. bulderi* exhibited the strongest acid-tolerant phenotype, maintaining growth at pH values as low as 1.5. Haploid *S. bombicola* also displayed pronounced acid tolerance, retaining growth at pH 1.5 at 30°C, whereas *S. batistae* showed robust growth down to pH 2.0 but failed to grow at pH 1.5. In comparison, the *S. cerevisiae* reference strains exhibited substantially reduced tolerance under strongly acidic conditions (Fig. 1). Although the particularly robust phenotype of *M. bulderi* may be associated with its diploid state, further investigation would be required to determine whether ploidy also contributes to acid tolerance.

Acid tolerance was further characterised by growth-curve analysis in liquid medium at 25°C and 30°C across decreasing pH conditions (Supplementary Fig. 1). At 25°C, the *S. cerevisiae* reference strains failed to sustain growth at pH 2.5, whereas *M. bulderi* maintained robust growth. *S. batistae* exhibited a longer lag phase than *M. bulderi* but ultimately reached a higher biomass during stationary phase. Similar trends were observed at 30°C. At pH 2.5 and 2.0, *M. bulderi* displayed the most rapid growth and shortest lag phase. *S. bombicola* grew more slowly than *M. bulderi*, with a longer lag phase, but initiated growth earlier than *S. batistae*. Despite its extended lag phase, *S. batistae* subsequently showed substantial biomass accumulation and reached a higher stationary-phase biomass than *S. bombicola*. *S. bombicola* ultimately reached a stationary-phase biomass comparable to that of *M. bulderi*, despite its slower initial growth.

The differences between the NCYs became particularly pronounced at pH 1.8. *M. bulderi* retained the strongest growth phenotype, reaching an OD₆₀₀ of approximately 0.7, whereas *S. bombicola* showed limited but detectable growth, reaching an OD₆₀₀ of approximately 0.2. In contrast, *S. batistae* was unable to grow under these highly acidic conditions. Notably, the *S. cerevisiae* reference strains showed greater sensitivity to acidic conditions in liquid culture than on solid medium. While limited growth was detectable on acidic solid medium, the reference strains failed to sustain growth at pH 2.5 in liquid culture, suggesting that acid tolerance is influenced by the cultivation environment.

Collectively, these results demonstrate that the selected NCYs thrive across low-pH conditions that strongly restrict *S. cerevisiae*, although their growth dynamics and lower pH limits differ substantially between species. *M. bulderi* exhibited the most robust phenotype under extremely acidic conditions, while *S. bombicola* and *S. batistae* also retained considerable acid tolerance. The ability of these NCYs to maintain growth under acidic conditions supports their selection as promising microbial chassis for sustainable lactic acid production, where tolerance to increasing medium acidity is an important host characteristic.

### Development of an efficient chemical transformation platform enables plasmid delivery into genetically intractable non-conventional yeasts

A major initial challenge was the inability to introduce plasmid DNA efficiently into *Maudiozyma bulderi*, even when using a plasmid containing a suitable origin of replication. The Pan/ARS-based CRISPR/Cas9 plasmid pUDPzb_U1, carrying the *hph* hygromycin resistance marker (Jayaprakash, Barroso et al. 2023) was therefore used as a test plasmid to establish a robust transformation procedure. As plasmid delivery represents an essential prerequisite for subsequent genetic engineering, several transformation conditions were screened and optimised to overcome the limited genetic accessibility of these non-conventional yeasts (NCYs).

We optimised the chemical transformation protocol of Lodens and co-workers (Lodens, De Graeve et al. 2018), and successfully enabled plasmid delivery into *M. bulderi*, *Starmerella batistae*, *Starmerella bombicola*, *Kazachstania aerobia*, *Kazachstania naganishii*, and *Yarrowia lipolytica* (Fig. 2). Notably, our optimised protocol (see material and methods) was also effective in *Y. lipolytica*, a species for which dedicated transformation procedures are already established, further demonstrating the cross-species applicability of the optimised method. Five independent transformants from each species were analysed by colony PCR, confirming successful plasmid acquisition with a high proportion of PCR-positive colonies.

**Figure 2.**
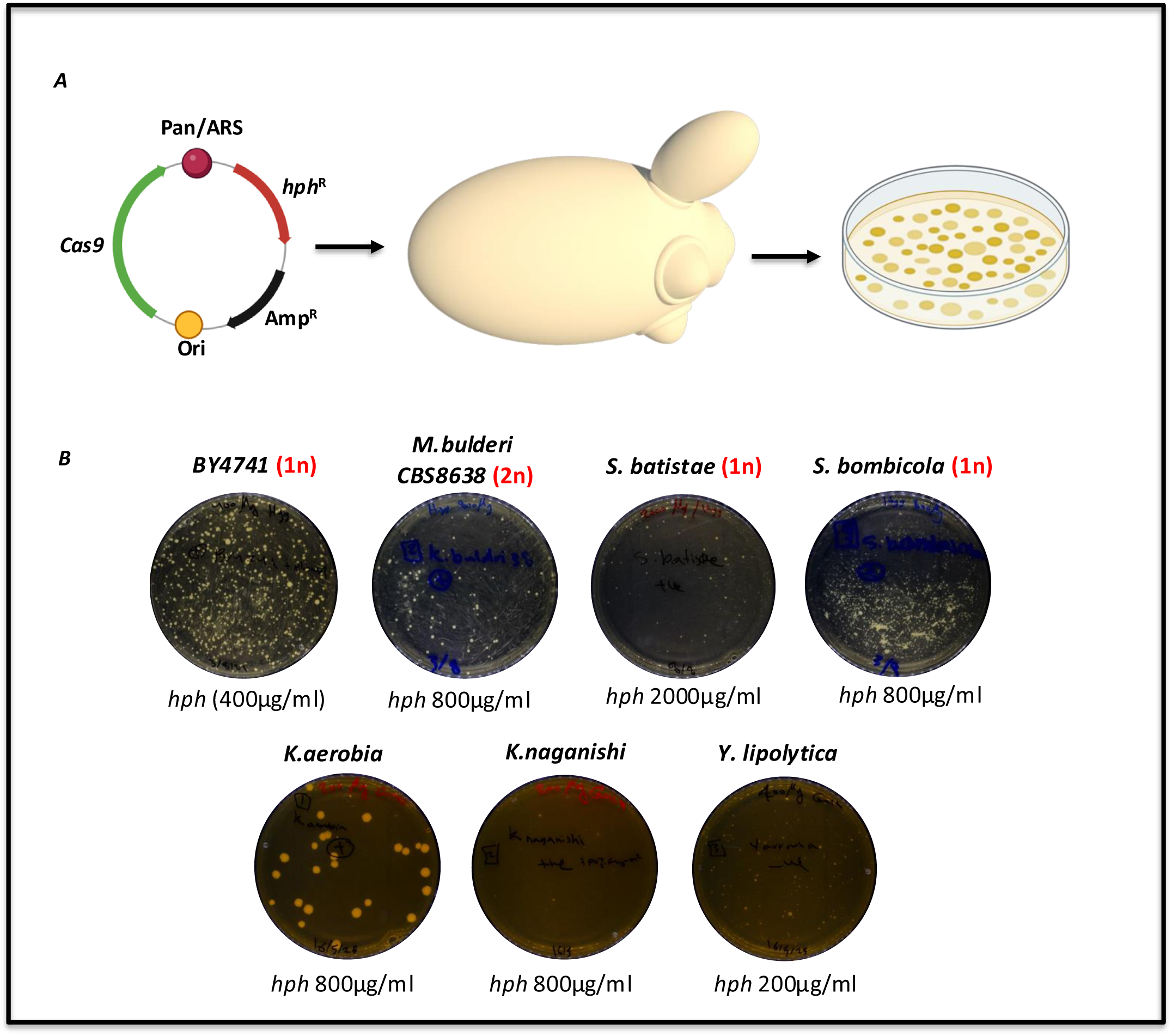
Development of an efficient chemical transformation platform enables plasmid delivery into genetically intractable non-conventional yeasts. **(A)** Schematic overview of the optimised chemical transformation workflow used for plasmid delivery into non-conventional yeasts (NCYs). The Pan/ARS-based pUDPzb_U1 carrying the *hph* hygromycin resistance marker (Jayaprakash, Barroso et al. 2023) was used as the test plasmid. **(B)** Representative transformants obtained following chemical transformation. *Saccharomyces cerevisiae* was used as the positive control and selected on YPD supplemented with 100 μg mL⁻¹ hygromycin B, whereas non-conventional yeasts were selected on YPD containing 1,000 μg mL⁻¹ hygromycin. The hygromycin B concentrations used for each strain are indicated below the corresponding plates.

Although successful transformation was achieved across all tested NCYs, the number of transformants recovered was generally lower than that obtained for *Saccharomyces cerevisiae*, which was included as a positive control. Moreover, approximately tenfold more plasmid DNA was required for NCY transformation than for *S. cerevisiae*. Species-specific differences in transformant morphology were also observed. In particular, *S. batistae* transformants frequently appeared as small colonies; nevertheless, the colonies analysed by PCR were confirmed to contain the plasmid.

A further challenge was the emergence of background colonies on negative-control plates lacking plasmid DNA. These colonies typically appeared following prolonged incubation (4–5 days), suggesting the emergence of spontaneous resistance under selective pressure. In contrast, genuine transformants generally appeared earlier, highlighting the importance of colony appearance time combined with PCR verification for reliable identification of positive transformants.

Collectively, optimisation of this chemical transformation procedure overcame a major barrier to the genetic manipulation of *M. bulderi* and established a single transformation approach applicable across phylogenetically diverse NCY species. This broad applicability provided the experimental foundation required for subsequent development of synthetic biology tools for heterologous gene expression and genome engineering.

### Development of a versatile synthetic biology toolkit enables heterologous gene expression and genome editing in non-conventional yeasts

Genetic engineering of non-conventional yeasts (NCYs) requires molecular tools compatible with their distinct replication and selection requirements. Conventional *Saccharomyces cerevisiae* plasmids cannot necessarily be maintained across phylogenetically diverse NCYs because episomal propagation requires a compatible origin of replication. Furthermore, efficient genome engineering of genetically less tractable species benefits from flexible CRISPR/Cas9 systems capable of accommodating multiple guide RNAs (gRNAs). To address these limitations, we developed two complementary Pan/ARS-based shuttle vector platforms: a modular YeastFab-compatible expression vector for heterologous gene and pathway expression and a CRISPR/Cas9 CREEPY vector supporting single and multiplex gRNA assembly (Fig. 3A–C). Detailed vector construction and cloning procedures are provided in Materials and Methods.

**Figure 3.**
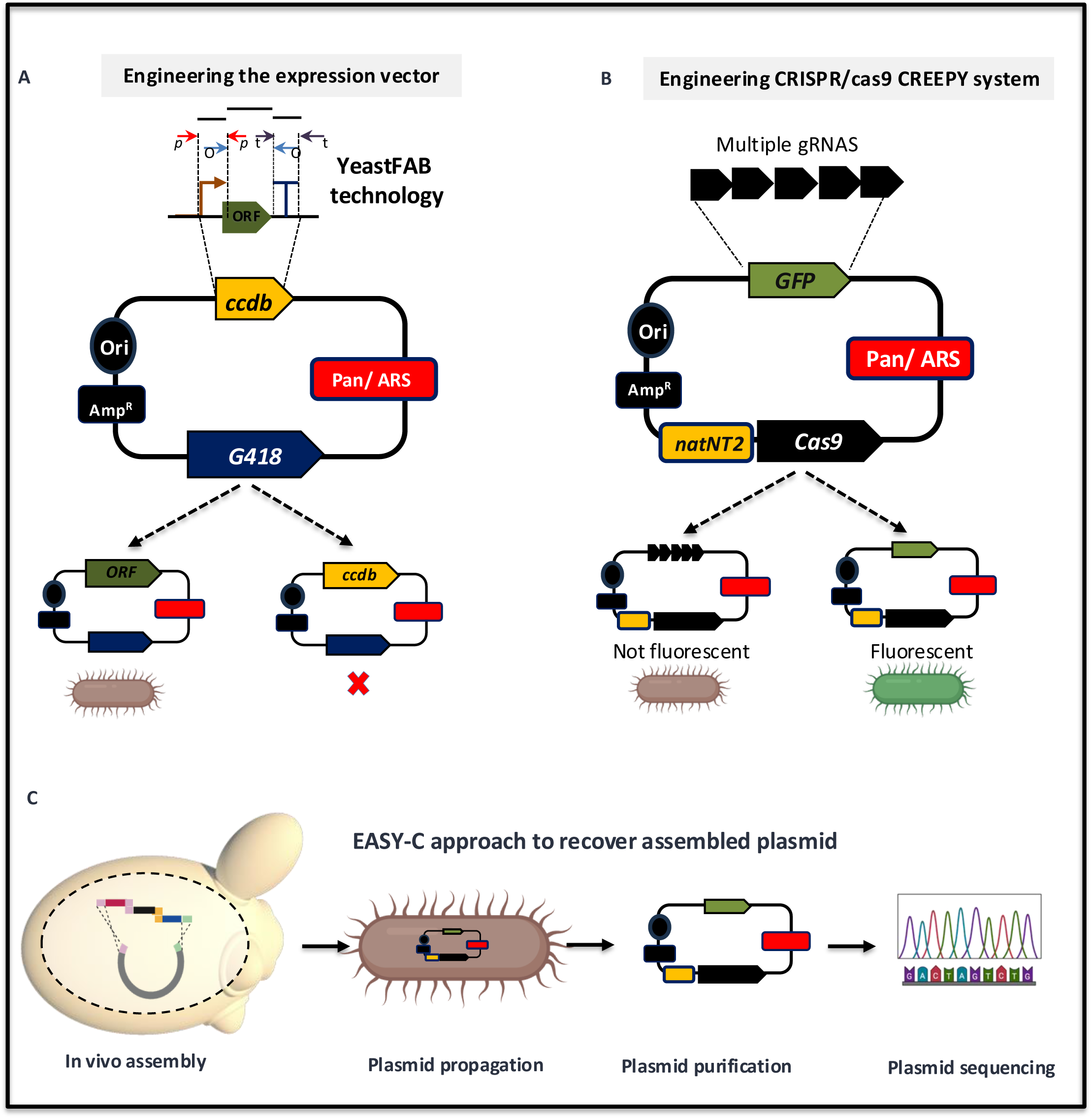
Development of a versatile synthetic biology toolkit enables heterologous gene expression and efficient genome editing in non-conventional yeasts. **(A)** Schematic representation of the broad-host-range Pan/ARS expression vector engineered as a shuttle plasmid for propagation in both non-conventional yeasts (NCYs) and *Escherichia coli*. The vector contains a Pan/ARS replication sequence enabling episomal maintenance in NCYs, a *kanMX* cassette conferring G418 resistance for yeast selection, an AmpR/ColE1 module for propagation and selection in *E. coli*, and a YeastFab-compatible *ccdB* destination cassette flanked by *BsmBI* recognition sites. The latter enables one-pot Golden Gate assembly of standardised promoter, open reading frame (ORF), and terminator modules. Recombinant constructs were initially identified by *ccdB* counter-selection and subsequently verified by restriction digestion and sequencing**. (B)** Schematic representation of the broad-host-range CRISPR/Cas9 CREEPY vector engineered for propagation in NCYs and *E. coli*. The vector contains a Pan/ARS replication sequence, a *natNT2* selectable marker for yeast, and an AmpR/ColE1 module for bacterial propagation. A superfolder GFP (*sfGFP*) cassette flanked by *BsmBI* recognition sites enables one-pot Golden Gate assembly of single or multiplex guide RNA (gRNA) expression cassettes. Successful gRNA assembly is identified by loss of GFP fluorescence following replacement of the *sfGFP* cassette. **(C)** In vivo assembly and recovery workflow for the Pan/ARS expression and CRISPR/Cas9 CREEPY vectors. Both vectors were assembled in *Saccharomyces cerevisiae* using its homologous recombination machinery. Correct assembly was confirmed by junction PCR, followed by plasmid recovery using the EASY-C workflow, propagation in *E. coli* DH5α, and full-length plasmid sequence verification by Nanopore sequencing.

The compact (∼6 kb) broad-host-range expression vector incorporated the pan-fungal autonomously replicating sequence (Pan/ARS) to support episomal maintenance across conventional and non-conventional yeasts (Jayaprakash, Barroso et al. 2023), *kanMX* for G418 selection in yeast, and an AmpR/ColE1 module for propagation and selection in *Escherichia coli* (Fig. 3A). To enable standardised assembly of heterologous expression cassettes, a *ccdB*-based destination cassette flanked by Type IIS restriction sites was incorporated according to the YeastFab framework (Guo, Dong et al. 2015, Garcia-Ruiz, Auxillos et al. 2018). This architecture enables one-pot Golden Gate assembly of promoter, open reading frame (ORF), and terminator modules, providing a reusable platform for construction of individual transcriptional units and multigene pathways.

In parallel, a broad-host-range CRISPR/Cas9 shuttle vector was developed (Fig. 3B) by adapting the CREEPY genome-editing system (Zhao, Coelho et al. 2023). The vector combined Pan/ARS-mediated replication with a *Cas9* expression cassette, *natNT2* for nourseothricin selection in yeast, and an AmpR/ColE1 module for propagation in *E. coli*. A superfolder GFP (*sfGFP*) cloning cassette (Pedelacq, Cabantous et al. 2006) , flanked by *BsmBI* recognition sites, was incorporated to enable one-pot Golden Gate assembly of single or multiplex gRNA expression cassettes. Replacement of the *sfGFP* cassette by the gRNA module results in loss of green fluorescence, providing a rapid visual screen for successful assembly. Importantly, the modular architecture enables multiple gRNAs to be incorporated within a single construct, allowing simultaneous targeting of multiple positions within a genomic locus and providing a flexible strategy for genome editing in genetically less tractable NCYs.

Both vectors were assembled using a common *S. cerevisiae*-based in vivo homologous-recombination workflow (Fig. 3C). This strategy proved particularly useful for complex multi-fragment assemblies that had been challenging to generate using conventional in vitro cloning during vector development. Correct assemblies were identified by junction PCR and subsequently recovered directly from yeast using the EASY-C workflow(Swidah, Monti et al. 2026). Because the expression vector contains the *ccdB* counter-selection cassette, the recovered expression plasmid was propagated in *ccdB*-resistant *E. coli*, whereas the recovered CRISPR/Cas9 plasmid was propagated in standard *E. coli* DH5α. The recovered vectors were subsequently purified, and sequence verified. For the CRISPR/Cas9 vector, an internal *BsmBI* recognition site within the *Cas9* coding sequence was additionally removed by synonymous mutagenesis to ensure compatibility with Golden Gate cloning.

Coupling *S. cerevisiae*-mediated homologous recombination with EASY-C recovery therefore provided a streamlined workflow for the construction, recovery, and validation of complex multi-fragment vectors, overcoming difficulties encountered with conventional in vitro assembly during vector development. The resulting Pan/ARS expression and CRISPR/Cas9 CREEPY vectors establish a complementary synthetic biology toolkit combining broad-host-range heterologous gene expression with single-and multiplex-gRNA genome editing in NCYs (Fig. 3). These tools provided the foundation for subsequent heterologous lactate dehydrogenase expression and targeted metabolic engineering of the selected acidophilic NCY chassis.

### YeastFab enables refactoring of regulatory elements for bacterial lactate dehydrogenase gene expression in non-conventional yeasts

Following the development of the broad-host-range Pan/ARS expression platform, we applied the YeastFab framework to introduce heterologous lactate dehydrogenase (*ldh*) genes into the selected acidophilic non-conventional yeast (NCY) chassis. A two-stage engineering strategy was established, comprising heterologous *ldh* expression followed by species-specific modification of competing carbon metabolism (Fig. 4A).

**Figure 4.**
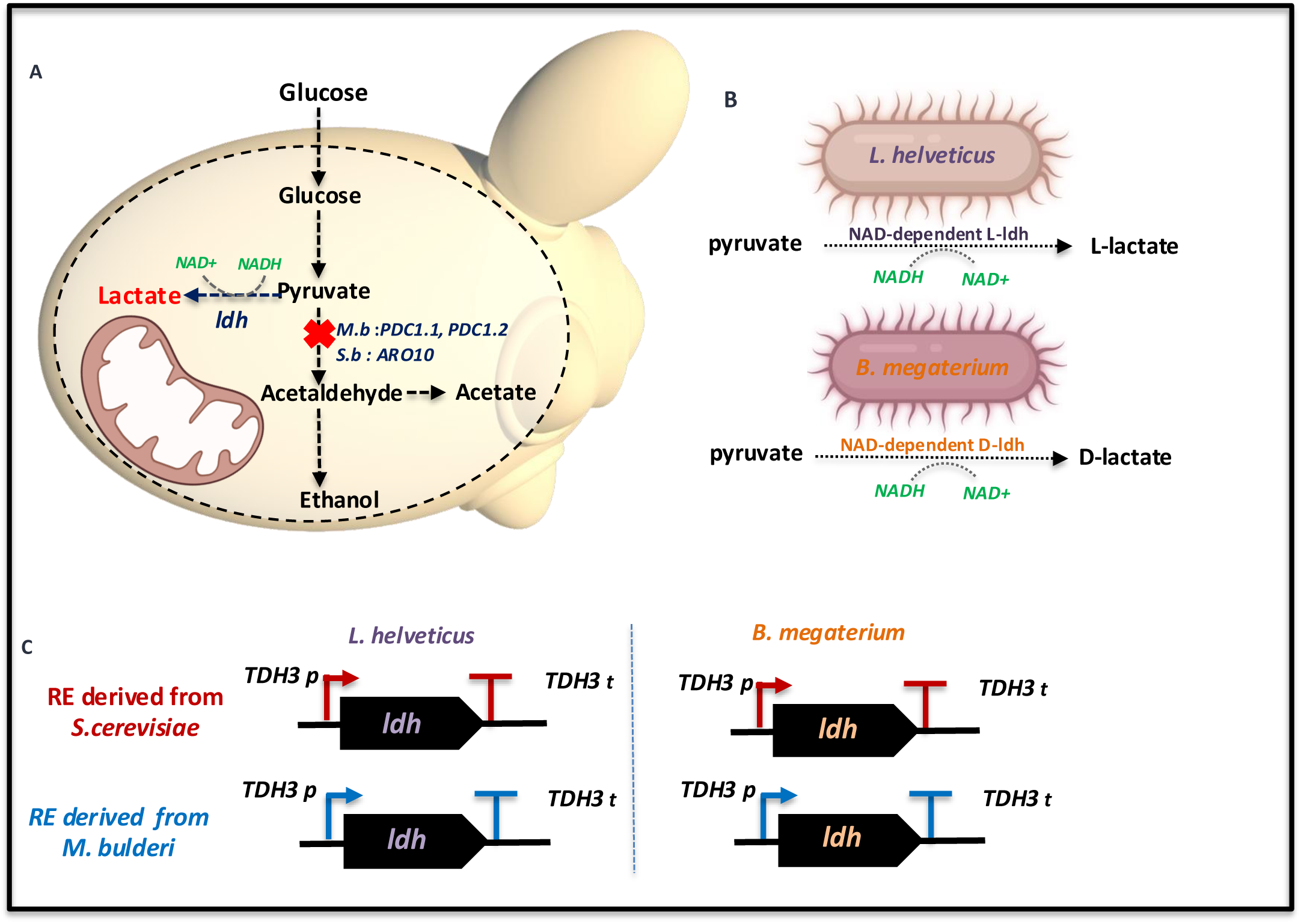
YeastFab-mediated assembly of bacterial lactate dehydrogenase expression constructs and metabolic engineering strategy for non-conventional yeasts. **(A)** Schematic representation of the two-stage engineering strategy designed to improve lactic acid production in acid-tolerant non-conventional yeasts. In the first stage, heterologous *ldh* expression was introduced to enhance the conversion of pyruvate to lactate using NADH as a cofactor (green). In the second stage, competing pyruvate-consuming pathways were identified as targets for metabolic engineering, including the pyruvate decarboxylase genes *PDC1.1* and *PDC1.2* in *Maudiozyma bulderi* and the phenylpyruvate decarboxylase gene *ARO10* in *Starmerella* spp. Targeted pathway engineering was successfully demonstrated through *ARO10* deletion in *S. bombicola*. **(B)** Schematic representation of the two heterologous *ldh* genes selected for expression in non-conventional yeasts. The *Lactobacillus helveticus ldh* encodes an NAD-dependent L-lactate dehydrogenase that catalyses the conversion of pyruvate to L-lactate (purple), whereas the *Bacillus megaterium ldh* encodes an NAD-dependent D-lactate dehydrogenase that catalyses the conversion of pyruvate to D-lactate (orange).

In the first stage, two bacterial *ldh* genes with different stereospecificities were selected (Fig. 4B): *Lactobacillus helveticus ldh*, encoding an NAD-dependent L-lactate dehydrogenase, and *Bacillus megaterium ldh*, encoding an NAD-dependent D-lactate dehydrogenase. Each *ldh* gene was assembled with *TDH3* regulatory elements derived from either *S. cerevisiae* or *M. bulderi* and introduced into the Pan/ARS broad-host-range expression vector (Fig. 4C). This generated four constructs for heterologous expression in the selected NCY chassis. Construct functionality was subsequently assessed by measuring lactic acid production using HPLC.

In the second stage, the competing pathways were targeted. In *M. bulderi*, the pyruvate decarboxylase genes *PDC1.1* and *PDC1.2* (Balarezo-Cisneros, Hanak et al. 2025), which encode enzymes involved in the decarboxylation of pyruvate to acetaldehyde, were selected. In *S. bombicola*, the corresponding target was *ARO10*, which encodes a putative broad-substrate-specificity 2-oxo-acid decarboxylase involved in the competing Ehrlich pathway. *ARO10* function has been characterised in *S. cerevisiae*, and homologues have also been identified in non-conventional yeasts, including *Yarrowia lipolytica* (Vuralhan, Morais et al. 2003, Vuralhan, Luttik et al. 2005, Celinska, Borkowska et al. 2019).

*ARO10* deletion was successfully achieved in *S. bombicola*, enabling subsequent evaluation of the combined effects of endogenous pathway engineering and heterologous *ldh* expression on lactic acid production. The deletion cassette was integrated into the genome of *M. bulderi*, however, it was off-target in the transformants tested. Extension of the *ARO10* deletion strategy to *S. batistae* was not pursued within the timeframe of this study.

### Pan/ARS-*ldh* expression vectors are stably maintained in non-conventional yeasts over 100 generations

Following engineering of the Pan/ARS-*ldh* expression vectors, we next investigated whether these constructs could be stably maintained during prolonged cultivation. *M. bulderi*, *S. bombicola*, and *S. batistae* strains harbouring each of the four *ldh* expression constructs, together with their corresponding empty-vector controls, were serially propagated for 100 generations under G418 selection (Fig. S2A). Following 100 generations, plasmid retention was assessed by colony PCR using five independent colonies for each construct. Expected amplification products were detected across the constructs analysed in all three NCY species, demonstrating robust maintenance of the Pan/ARS-*ldh* expression vectors during prolonged cultivation under selective conditions. Representative colony PCR analysis of *S. bombicola* harbouring the four *ldh* constructs and the empty Pan/ARS vector is shown in Fig. S2A.

Together, these results demonstrate that the Pan/ARS-YeastFab platform supports long-term maintenance of heterologous *ldh* expression vectors across multiple acid-tolerant NCYs. Having established stable plasmid maintenance, we next investigated the functional consequences of heterologous *ldh* expression on lactic acid production and its interaction with endogenous lactic acid production in the different NCY chassis.

### Non-conventional yeasts possess endogenous lactic acid production and exhibit species-specific responses to heterologous lactate dehydrogenase expression

We evaluated lactic acid production after 7 days of cultivation at 30°C in YPD supplemented with G418 (YPD-G418) to maintain selection for the Pan/ARS expression plasmids. Notably, all three non-conventional yeast species produced lactic acid in the absence of heterologous *ldh* expression, revealing endogenous lactic acid production under the conditions tested.

*Starmerella batistae* harbouring the empty Pan/ARS vector produced approximately 3 g L⁻¹ lactic acid from 2% glucose (Fig. 5A). Heterologous *ldh* expression further enhanced production in a construct-dependent manner. The highest titre, approximately 6 g L⁻¹, was obtained with the *Bacillus megaterium ldh* expressed using *Saccharomyces cerevisiae*-derived regulatory elements, representing an approximately twofold increase relative to the empty-vector control. Biomass-normalised analysis showed a similar pattern, with the *B. megaterium ldh*/*S. cerevisiae* regulatory-element combination providing the clearest enhancement in lactic acid production per cell. A more modest increase, to approximately 4 g L⁻¹, was observed with the *Lactobacillus helveticus ldh* expressed using *S. cerevisiae*-derived regulatory elements. However, this improvement was less apparent following normalisation to OD₆₀₀ and was not statistically significant, indicating that the increase in titre was at least partly associated with differences in biomass accumulation (Supplementary Fig. 3A).

**Figure 5.**
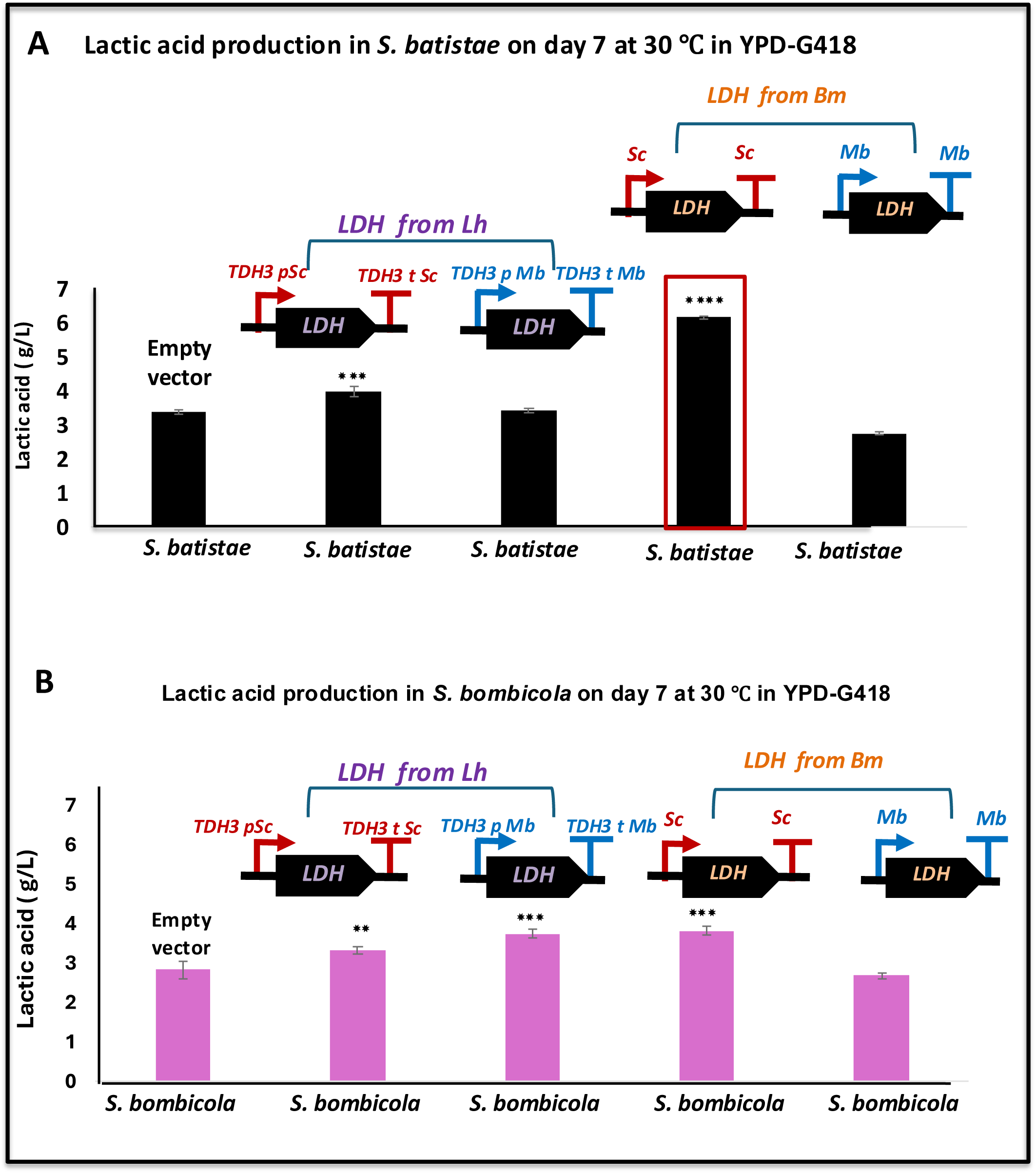
Non-conventional yeasts possess endogenous lactic acid biosynthetic pathways with species-specific responses to heterologous lactate dehydrogenase expression. **(A)** Lactic acid production by *Starmerella batistae* harbouring the empty Pan/ARS vector or four heterologous *ldh* expression constructs**. (B)** Lactic acid production by *Starmerella bombicola* harbouring the corresponding empty Pan/ARS vector or four heterologous *ldh* expression constructs. The constructs contained *ldh* derived from *Lactobacillus helveticus* (L-LDH; purple) or *Bacillus megaterium* (D-LDH; orange), expressed using *TDH3* regulatory elements derived from either *Saccharomyces cerevisiae* (red) or *Maudiozyma bulderi* (blue), as indicated above the corresponding bars. Empty-vector controls are shown in black. Cultures were incubated at 30°C, and samples were collected on day 7 for lactic acid quantification by HPLC. Data represent the mean ± standard deviation (SD) of three independent biological replicates. Statistical significance was assessed by one-way ANOVA followed by Dunnett’s multiple-comparisons test, comparing each *ldh* construct with the corresponding empty-vector control. Significance is indicated as ns, *P* ≥ 0.05; *, *P* < 0.05; **, *P* < 0.01; ***, *P* < 0.001; and ****, *P* < 0.0001.

Endogenous lactic acid production was also observed in *Starmerella bombicola*, with the empty-vector strain producing approximately 3 g L⁻¹ lactic acid (Fig. 5B). In contrast to *S. batistae*, the effects of heterologous *ldh* expression were more modest. Increased lactic acid titres were observed with *L. helveticus ldh* expressed using *M. bulderi*-derived regulatory elements and with *B. megaterium ldh* expressed using *S. cerevisiae*-derived regulatory elements, reaching approximately 3.5–4 g L⁻¹. These differences were also evident following normalisation to biomass (Supplementary Fig. 3B), further demonstrating that the response to heterologous LDH expression depends on both the LDH enzyme and its regulatory context.

*Maudiozyma bulderi* displayed a slow-growing phenotype and produced endogenous lactic acid, with the empty-vector strain producing approximately 3 g L⁻¹ lactic acid). Unlike the two *Starmerella* species tested, heterologous *ldh* expression did not increase the final lactic acid titre and was associated with reduced growth and biomass accumulation (data not shown). Successful heterologous LDH protein expression was nevertheless confirmed by Western blotting (Fig. 4D), indicating that the absence of increased lactic acid production was not simply attributable to failure of protein expression. Instead, the reduced growth associated with *ldh* expression suggests that heterologous LDH activity imposes a fitness burden in *M. bulderi*. This phenotype may reflect metabolic or redox constraints associated with increased NADH-dependent conversion of pyruvate to lactate, although further investigation is required to establish the underlying mechanism. Further optimisation of the expression system, including modulation of plasmid copy number and promoter strength and optimisation of *ldh* codon usage, may therefore be required to balance heterologous LDH expression with host fitness and improve lactic acid production in *M. bulderi*.

These results demonstrate endogenous lactic acid production across the three acid-tolerant NCYs investigated and reveal strongly species-and construct-dependent responses to heterologous *ldh* expression. Whereas *S. batistae* showed a substantial enhancement in lactic acid production with an appropriate *ldh* construct, *S. bombicola* displayed more moderate improvements, while *M. bulderi* exhibited an apparent fitness cost associated with heterologous expression. These contrasting responses highlight the importance of balancing LDH activity with host physiology and tailoring enzyme origin and regulatory architecture to individual NCY chassis for improved lactic acid production.

### Deletion of *ARO10* redirects carbon flux and enhances endogenous and heterologous lactic acid production in *Starmerella bombicola*

Given the endogenous lactic acid production observed in the NCYs, we next investigated whether metabolic engineering could further enhance production by redirecting carbon flux towards lactate. *ARO10*, encoding a phenylpyruvate decarboxylase involved in the decarboxylation of aromatic and branched-chain α-keto acids, was selected as a metabolic engineering target in *S. bombicola*. We hypothesised that disruption of this competing decarboxylation route could increase carbon availability for lactate formation.

*ARO10* was deleted using a *natNT2* deletion cassette flanked by approximately 800-bp homology regions, combined with the broad-host-range CRISPR/Cas9 CREEPY system developed in this study. Three gRNAs targeting the beginning, middle and end of the *ARO10* ORF were used to promote efficient genome editing. Screening of 96 transformants by diagnostic PCR demonstrated a deletion efficiency of >94%, highlighting the high efficiency of the multiplex CRISPR/Cas9 strategy for genome engineering in *S. bombicola*.

Deletion of *ARO10* alone increased endogenous lactic acid production from ca. 3 g L⁻¹ in the parental strain to ca. 4 g L⁻¹ in the Δ*ARO10* strain (Fig. 6A). Strains were cultivated at 30°C in YPD, and samples were collected on day 7 for HPLC analysis. The increase was also retained following normalisation to OD₆₀₀ (Supplementary Fig. 4A), indicating that the enhanced titre was not solely attributable to differences in biomass accumulation and supporting increased lactic acid production at the cellular level.

**Figure 6.**
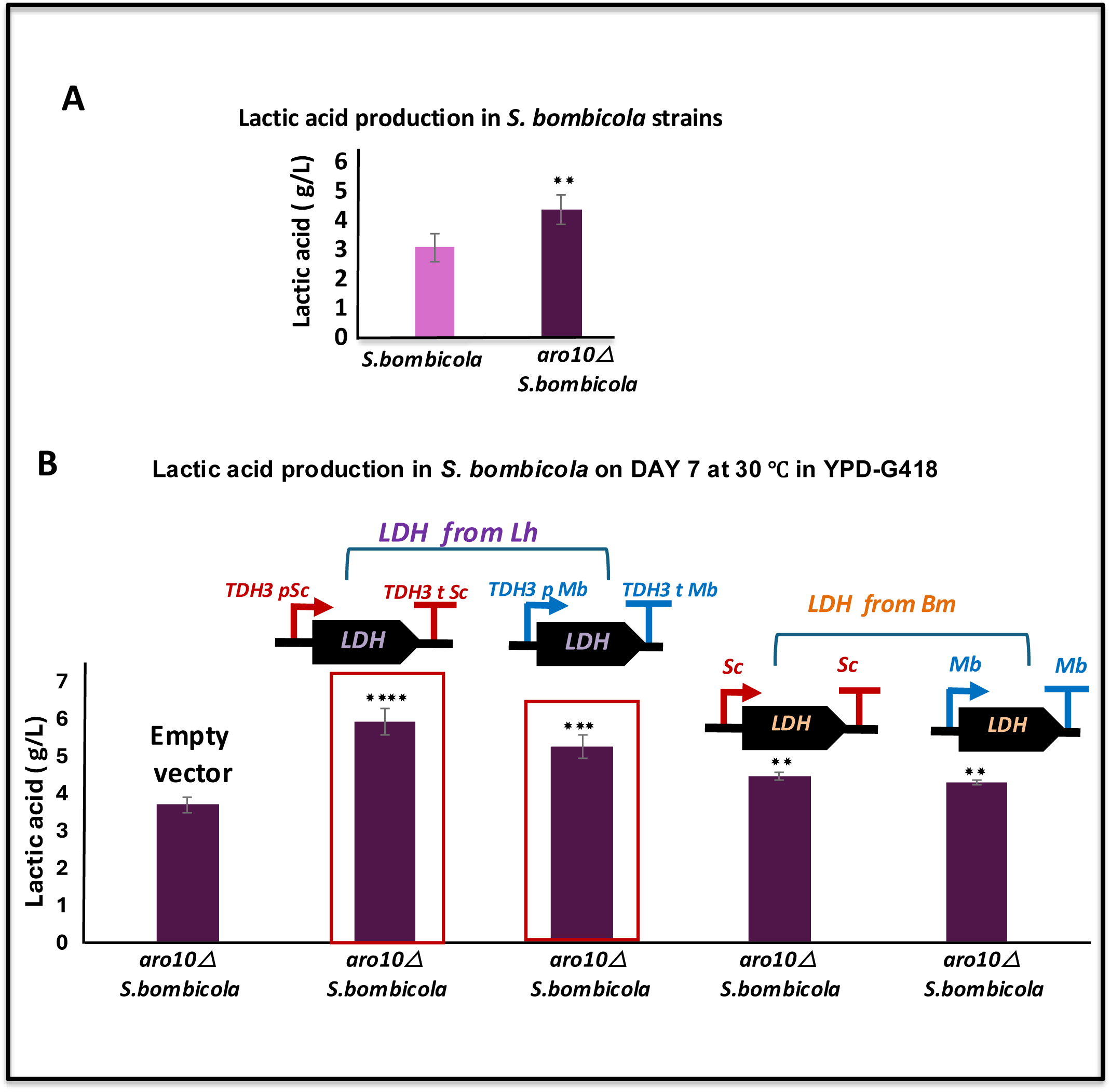
Deletion of *ARO10* redirects carbon flux and enhances endogenous and heterologous lactic acid production in *Starmerella bombicola* (A) Effect of *ARO10* deletion on endogenous lactic acid production in *S. bombicola*, showing increased lactic acid production following *ARO10* deletion in the absence of heterologous *ldh* expression. Statistical significance was assessed using a two-tailed unpaired Student’s *t*-test (*P < 0.01).* (B) Lactic acid production by *Δ*ARO10 S. bombicola* harbouring the empty Pan/ARS vector or four heterologous *ldh* expression constructs. The constructs contained *ldh* derived from *Lactobacillus helveticus* (L-LDH; purple) or *Bacillus megaterium* (D-LDH; orange), expressed using *TDH3* regulatory elements derived from either *Saccharomyces cerevisiae* (red) or *Maudiozyma bulderi* (blue), as indicated above the corresponding bars. Empty-vector controls are shown in black. Cultures were incubated at 30°C, and samples were collected on day 7 for lactic acid quantification by HPLC. Data represent the mean ± standard deviation (SD) of three independent biological replicates. Statistical significance in panel B was assessed by one-way ANOVA followed by Dunnett’s multiple-comparisons test, with each *ldh* construct compared with the Δ*ARO10* empty-vector control. Significance is indicated as ns, *P* ≥ 0.05; *, *P* < 0.05; **, *P* < 0.01; ***, *P* < 0.001; and ****, *P* < 0.0001.

We next investigated whether *ARO10* deletion could be combined with heterologous *ldh* expression to further enhance lactic acid production. Strains were cultivated at 30°C in YPD-G418, and samples were collected on day 7 for HPLC analysis. Notably, expression of the *Lactobacillus helveticus ldh* using *S. cerevisiae*- or *M. bulderi*-derived regulatory elements in the Δ*ARO10* background increased lactic acid titres to approximately 6 and 5 g L⁻¹, respectively (Fig. 6B). These results demonstrate that metabolic redirection through *ARO10* deletion can be combined with heterologous LDH activity to further improve lactic acid production. Biomass-normalised analysis showed a corresponding improvement, particularly for the *L. helveticus ldh* constructs (Supplementary Fig. 4B).

These results identify *ARO10* as an effective metabolic engineering target in *S. bombicola*, with its deletion enhancing both endogenous and heterologous lactic acid production.

Together, these findings establish a proof-of-concept strategy for combining endogenous pathway engineering with heterologous *ldh* expression and provide a framework for extending this approach to additional acid-tolerant NCY chassis.

### Non-conventional yeasts display distinct capacities for L- and D-lactate utilisation

An important consideration when developing yeasts for lactic acid production is whether the host can reutilise the accumulated product as a carbon source, potentially reducing the final product titre. We therefore assessed the capacity of *M. bulderi*, *S. batistae*, and *S. bombicola* to utilise L- and D-lactate. Growth was monitored for 3 days at 30°C in media containing decreasing concentrations of glucose supplemented with increasing concentrations of either L- or D-lactate: 2% glucose, 1.5% glucose + 0.5% lactate, 1% glucose + 1% lactate, or 2% lactate as the sole carbon source (Fig. 7).

**Figure 7.**
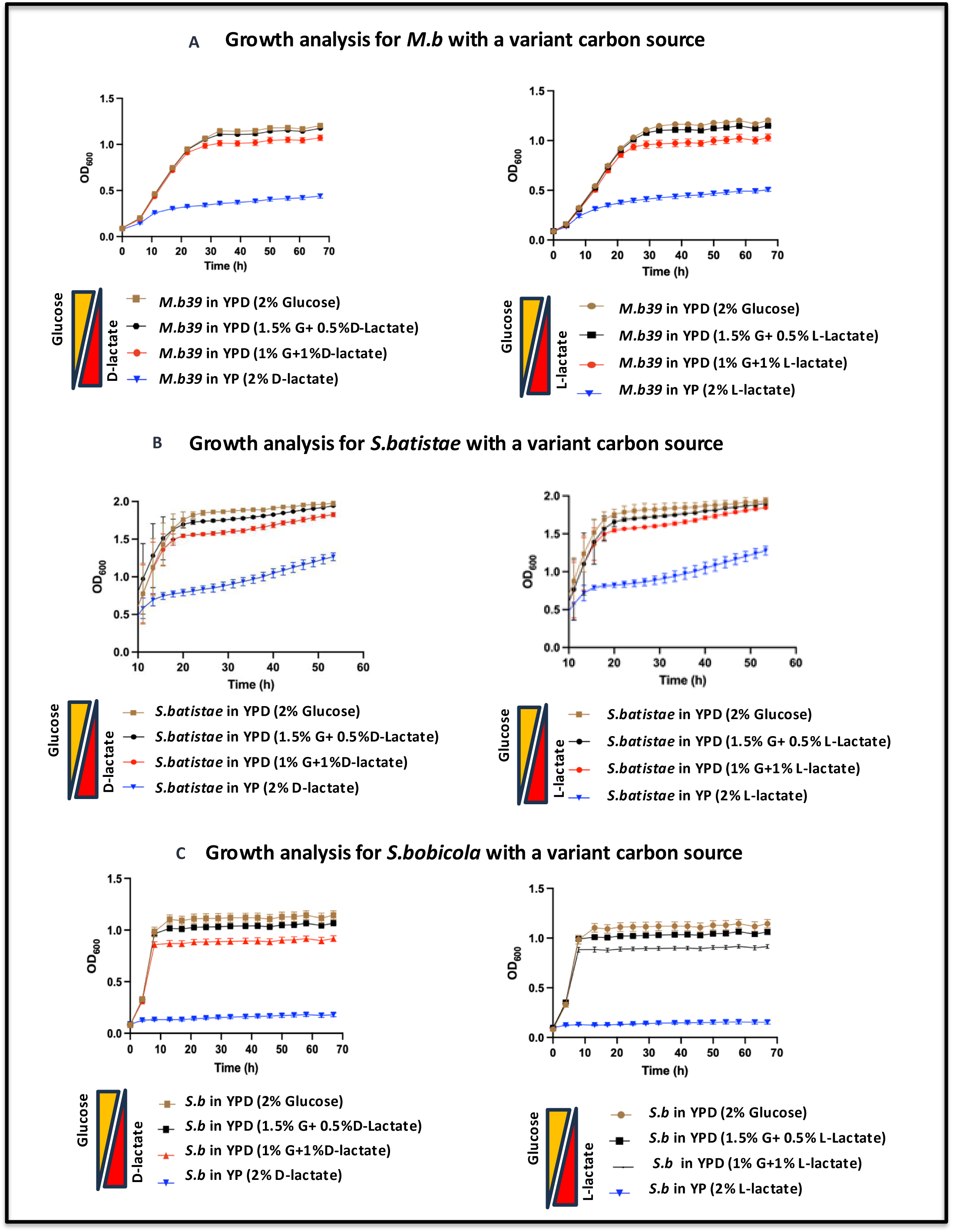
Non-conventional yeasts display distinct capacities for L- and D-lactate utilisation. Growth-curve analysis of *Maudiozyma bulderi* **(A)**, *Starmerella batistae* **(B)**, and *Starmerella bombicola* **(C)** during cultivation at 30°C for 3 days in media containing glucose, L-lactate, or D-lactate as carbon sources. For each species, the left panel shows growth in 2% glucose, 1.5% glucose + 0.5% L-lactate, 1% glucose + 1% L-lactate, or 2% L-lactate, whereas the right panel shows the corresponding conditions containing D-lactate. Data represent the mean ± standard deviation (SD) of three independent biological replicates.

Across the three species, the highest biomass accumulation was generally observed with 2% glucose as the sole carbon source. Growth progressively decreased as the proportion of glucose was reduced and replaced with either L- or D-lactate, although the magnitude of this response differed substantially among species. *M. bulderi* displayed a limited but detectable capacity to utilise lactate, reaching an OD₆₀₀ of approximately 0.5 when either L- or D-lactate was provided as the sole carbon source (Fig. 7A). In contrast, *S. batistae* exhibited a pronounced capacity for lactate utilisation, reaching an OD₆₀₀ >1 with either L- or D-lactate as the sole carbon source (Fig. 7B). These results demonstrate that *S. batistae* can efficiently utilise both lactate stereoisomers in the absence of glucose.

*S. bombicola* showed little to no growth when either L- or D-lactate was supplied as the sole carbon source (Fig. 7C), indicating a limited capacity to reutilise extracellular lactate under the conditions tested. This phenotype represents a potentially advantageous characteristic for lactic acid bioproduction, as reduced product reutilisation could favour extracellular lactate accumulation compared with hosts such as *M. bulderi* and *S. batistae* that can utilise lactate as a carbon source.

Together, these results reveal marked species-specific differences in L- and D-lactate utilisation among the three acid-tolerant NCYs. In particular, the combination of endogenous lactic acid production, responsiveness to metabolic engineering, and limited lactate reutilisation further supports *S. bombicola* as a promising chassis for future development of lactic acid-producing yeast cell factories.

## Discussion

Acid-tolerant yeasts therefore represent attractive alternative production hosts, as their ability to grow at low pH could reduce the need for neutralisation. Nevertheless, acid tolerance alone is insufficient to establish an efficient low-pH fermentation process, and suitable hosts must combine robust growth with appropriate metabolic and physiological characteristics. *M. bulderi* grows under highly acidic conditions and tolerates elevated lactic acid concentrations (Balarezo-Cisneros, Hanak et al. 2025), while *S. batistae* has been reported to grow at pH 2 and in the presence of lactic acid (Timouma, Hanak et al. 2026). Together with the low-pH growth observed here for *S. bombicola*, these properties motivated our investigation of these three species as potential hosts for lactic acid production.

The domestication of these non-conventional yeasts (NCYs) requires genetic tools that function efficiently across species. Replication origins used in conventional *Saccharomyces cerevisiae* plasmids may not support plasmid maintenance in phylogenetically distant yeasts (Liachko and Dunham 2014). Moreover, the limited availability of auxotrophic strains often necessitates antibiotic-based selection, requiring species-specific optimisation of antibiotic concentrations and molecular confirmation of transformants (Liachko and Dunham 2014; Juergens, Varela et al. 2018). Targeted genome modification presents an additional challenge because non-homologous end joining can compete with homologous recombination in some NCYs (Juergens, Varela et al. 2018). To overcome these limitations, we adapted the chemical transformation method of Lodens, De Graeve et al. (2018) for *M. bulderi*, *S. bombicola*, *S. batistae*, *Kazachstania aerobia*, *K. naganishii* and *Yarrowia lipolytica*. The Pan/ARS-containing vector pUDBzB-41 provided a common platform for DNA delivery across these hosts (Jayaprakash, Barroso et al. 2023), and transformation conditions, including selective antibiotic concentrations, were optimised for each species. Together, these advances establish a platform for investigating native metabolism and developing engineered NCY production strains.

As proof of concept, we engineered NCYs for lactic acid production. Lactic acid is commonly produced using bacterial fermentation; however, many bacterial processes require continuous neutralisation as lactic acid accumulates, increasing downstream processing requirements and generating substantial salt waste (Abdel-Rahman, Tashiro et al. 2013; Juturu and Wu 2016).

We designed Pan/ARS expression vectors to enable systematic testing of heterologous genes and regulatory elements across different hosts. Compatibility with YeastFab allowed standardised promoters, open reading frames and terminators to be assembled in one-pot Golden Gate reactions. This modular architecture facilitated refactoring of codon-optimised ldh genes using native or orthogonal regulatory elements and enabled rapid evaluation of different expression configurations across yeast species (Guo, Dong et al. 2015; Garcia-Ruiz, Auxillos et al. 2018). When conventional cloning proved challenging, constructs could alternatively be generated by in vivo assembly in *S. cerevisiae*. Plasmid recovery using EASY-C subsequently enabled restriction-digest analysis and full-plasmid sequencing for construct verification (Swidah, Monti et al. 2026). This combination of modular assembly and sequence-level verification provides considerable flexibility for engineering underexplored yeasts. Importantly, Pan/ARS-ldh vectors were maintained in *M. bulderi*, S. *bombicola* and *S. batistae* for 100 generations under G418 selection, demonstrating stable maintenance during prolonged cultivation under selective conditions.

To complement the expression platform, we developed a modular CREEPY-derived CRISPR–Cas9 vector for genome engineering in NCYs. Starting from the backbone described by Jayaprakash, Barroso et al. (2023), we retained the Pan/ARS replication element to support plasmid maintenance across target hosts and incorporated natNT2 for selection. GFP was included to facilitate multiplex gRNA assembly and provide an initial screen for candidate constructs before sequence verification. By adapting the CREEPY strategy described by Zhao, Coelho et al. (2023), the resulting system combines broad-host plasmid replication with modular gRNA assembly and a convenient screening step. The effectiveness of this approach was demonstrated in *S. bombicola*, where deletion of *ARO10* was achieved at an efficiency exceeding 94%. Although editing performance remains to be established in the other NCYs examined here, these results demonstrate the potential of the system for efficient genome engineering in *S. bombicola*.

The three focal species showed marked differences in both lactic acid production and utilisation. Unengineered *M. bulderi*, *S. bombicola* and *S. batistae* all produced detectable quantities of lactic acid, indicating the presence of endogenous routes for lactate formation. However, their ability to utilise lactate differed substantially. When supplied as the sole carbon source, *S. bombicola* showed no detectable growth on either L- or D-lactate, *M. bulderi* displayed only limited growth, whereas S*. batistae* grew robustly on both forms. The inability of *S. bombicola* to readily utilise lactate could be advantageous for production by limiting product re-assimilation, whereas lactate utilisation by S*. batistae* could potentially reduce product accumulation during extended fermentation. These interpretations remain to be tested under production conditions, as growth on lactate as the sole carbon source does not establish whether lactate is consumed in the presence of glucose or following glucose depletion.

The effects of heterologous ldh expression were similarly host dependent, consistent with previous observations that yeast lactate production is strongly influenced by host metabolism, LDH origin and lactate export capacity (Branduardi, Sauer et al. 2006). *S. batistae* produced approximately 6 g/L lactic acid when the *B. megaterium* ldh gene was expressed using *S. cerevisiae* regulatory elements. In *S. bombicola*, deletion of *ARO10* enhanced lactic acid production both in the presence and absence of heterologous ldh expression, with an engineered strain also reaching approximately 6 g/L. As *ARO10* contributes to pyruvate decarboxylase activity and ethanol formation in *S. bombicola* (Goncalves, Wisecaver et al. 2018), increased lactate production following *ARO10* deletion is consistent with reduced competition for the pyruvate pool. However, measurements of ethanol, other fermentation products and carbon balance will be required to establish the underlying metabolic mechanism.

By contrast, introduction of an ldh expression vector into *M. bulderi* resulted in poor growth, suggesting a host-specific fitness cost associated with heterologous LDH expression. Disruption of cellular redox balance represents one possible explanation but will require direct assessment of growth, LDH activity, intracellular cofactors and metabolic products. More broadly, the native pathways responsible for lactic acid formation in all three species remain to be identified. Unengineered *S. cerevisiae*, for example, can generate D-lactate through methylglyoxal metabolism (Martins, Cordeiro et al. 2001; Stewart, Navid et al. 2013), and determining whether related or distinct pathways operate in these NCYs will be important for subsequent strain engineering.

The lactic acid titres of approximately 6 g/L achieved in this study therefore represent a proof of concept rather than an optimised production process. Higher titres have been achieved using engineered *Zygosaccharomyces bailii* and *Pichia kudriavzevii* under different strain and cultivation conditions (Kuanyshev, Rao et al. 2021; Zhang, Li et al. 2023), highlighting substantial scope for further improvement. A key priority will be to identify the native genes and metabolic routes responsible for endogenous lactate formation and determine how *ARO10* deletion and heterologous ldh expression affect ethanol formation, other by-products, growth and overall carbon allocation. Time-course experiments will also be required to establish whether accumulated lactate is consumed during glucose-containing fermentations or following glucose depletion.

Further optimisation should consider species-specific codon usage, LDH activity and the strength of native and orthogonal promoters and terminators. Our production experiments were performed in batch cultures; medium optimisation and comparison with fed-batch or continuous cultivation may identify conditions that better sustain biomass formation and lactic acid production. Separate quantification of L- and D-lactate and determination of optical purity will also be important. L-lactic acid is widely used in food and other applications and is the precursor of poly(L-lactic acid), whereas D-lactic acid can be used to produce poly (D-lactic acid); combining the two polymers can generate stereocomplex materials with enhanced thermal properties (Juturu and Wu 2016; Sun, Lu et al. 2022). The ability to direct production towards a specific lactate stereoisomer could therefore enable strains to be tailored to different industrial applications.

Ultimately, evaluation under controlled low-pH fermentation conditions will be required to establish titre, yield and productivity and to determine whether efficient production can be sustained without antibiotic selection. Nonetheless, the genetic tools established here substantially expand the engineering capabilities available for these underexplored acid-tolerant yeasts. By combining broad-host transformation and modular expression systems with efficient CRISPR–Cas9 genome editing, this study provides a foundation for developing NCYs as low-pH cell factories for the sustainable production of lactic acid and other bio-based chemicals.

## Author contributions

**D.D.** conceived the study**. R.S.** designed and engineered the genetic toolkit, optimised the chemical transformation protocol, constructed the plasmids, performed the experiments and HPLC analyses, analysed the data, and drafted the manuscript**. F.V.** and **M.H.G.** contributed to the HPLC analyses. **S.T.** performed computational analyses to identify candidate endogenous *LDH* genes in non-conventional yeasts. **F.V.** and **D.D.** acquired funding, supervised the research, contributed to scientific discussions, and critically revised the manuscript. All authors reviewed, edited, and approved the final manuscript.

## Supplementary data

All supplementary data are available in the Supplementary Information accompanying this article. Plasmid sequences and annotated vector maps will be deposited in Addgene and will be publicly available upon publication.

## Acknowledgements

We thank Professor Anthony Green, Director of the Manchester Institute of Biotechnology, for his support of R.S. and this research. We also thank members of the BP team for their critical discussions and constructive feedback during the study.

## Figure legends

Supplementary Figure 1. G**r**owth **curve analysis reveals distinct acidophilic profiles of non-conventional yeasts in liquid culture.** Growth of *Maudiozyma bulderi*, *Starmerella bombicola*, *Starmerella batistae*, and *Saccharomyces cerevisiae* reference strains was monitored in liquid medium across the indicated pH range at **25°C (A)** and **30°C (B)**. Growth was assessed by measuring OD₆₀₀ over time. Data represent the mean ± standard deviation (SD) of three independent biological replicates.

Supplementary Figure 2. P**l**asmid **stability of the broad-host expression vector harbouring *ldh* (A)** Schematic representation of the plasmid stability assay. Strains of *Maudiozyma bulderi*, *Starmerella bombicola*, and *Starmerella batistae* carrying the Pan/ARS-*ldh* expression plasmids or the corresponding empty Pan/ARS vector were serially propagated for 100 generations in liquid YPD supplemented with G418. Following cultivation, cells were plated onto selective YPD-G418 agar, and five independent biological colonies for each construct were analysed by colony PCR to assess plasmid retention. A representative colony PCR analysis of *S. bombicola* harbouring Pan/ARS-*ldh* constructs 1–4 and the empty Pan/ARS vector (construct 5) is shown on the right, demonstrating stable maintenance of the expression plasmids under selective conditions.

Supplementary Figure 3. Biomass-normalised lactic acid production reveals species-specific responses to heterologous *ldh* expression in non-conventional yeasts. Lactic acid production normalised to OD₆₀₀ in (A)*Maudiozyma bulderi* and (B) *Starmerella batistae* harbouring the empty Pan/ARS vector or heterologous *ldh* constructs containing regulatory elements derived from *Saccharomyces cerevisiae* or *M. bulderi*. Data represent the mean ± standard deviation (SD) of three independent biological replicates. Statistical significance was assessed by one-way ANOVA followed by Dunnett’s multiple-comparisons test, with each *ldh* construct compared with the empty-vector control. Significance is indicated as ns, *P* ≥ 0.05; *, *P* < 0.05; **, *P* < 0.01; ***, *P* < 0.001; and ****, *P* < 0.0001.

Supplementary Figure 4. Biomass-normalised lactic acid production reveals the impact of *ARO10* deletion on endogenous and heterologous lactic acid production in *Starmerella bombicola*. (A) Effect of *ARO10* deletion on endogenous lactic acid production in *Starmerella bombicola*, expressed as lactic acid production normalised to OD₆₀₀. Statistical significance was assessed using a two-tailed unpaired Student’s *t*-test (\**P* < 0.01). **(B)** Biomass-normalised lactic acid production in *S. bombicola* harbouring the empty Pan/ARS vector or four heterologous *ldh* expression constructs. The constructs contained *ldh* derived from *Lactobacillus helveticus* (L-LDH; purple) or *Bacillus megaterium* (D-LDH; orange), expressed using *TDH3* regulatory elements derived from either *Saccharomyces cerevisiae* (red) or *Maudiozyma bulderi* (blue), as indicated above the corresponding bars. Cultures were incubated at 30°C, and samples were collected on day 7 for lactic acid quantification by HPLC. Data represent the mean ± standard deviation (SD) of three independent biological replicates. Statistical significance in panel B was assessed by one-way ANOVA followed by Dunnett’s multiple-comparisons test, with each *ldh* construct compared with the corresponding empty-vector control. Significance is indicated as ns, *P* ≥ 0.05; *, *P* < 0.05; **, *P* < 0.01; ***, *P* < 0.001; and ****, *P* < 0.0001.

