## Supplementary Figures and Tables for "Engineering acidophilic non-conventional yeasts as low-pH cell factories for organic acid production"

**Supplementary Fig.1**

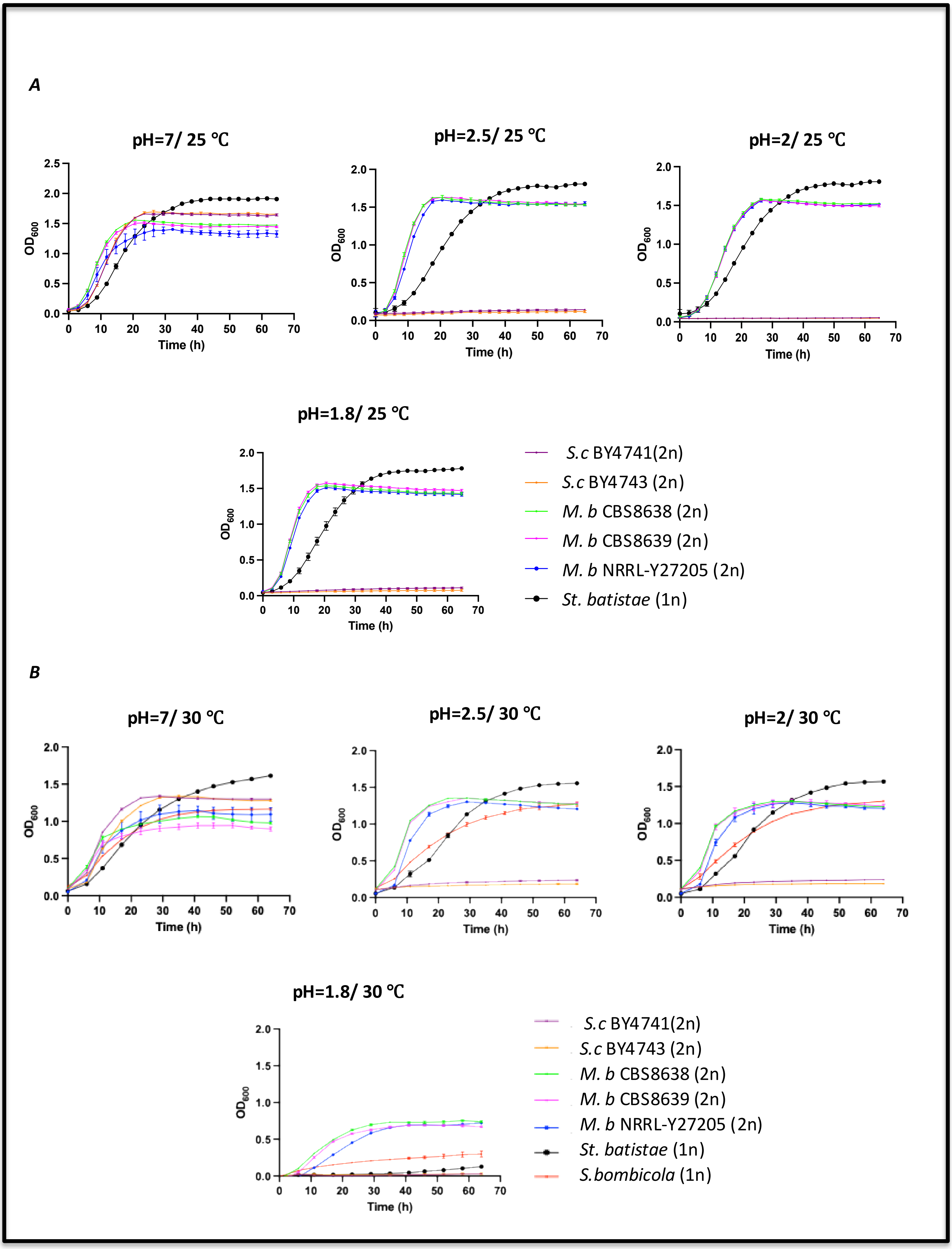


**Supplementary Figure 1. Growth curve analysis reveals distinct acidophilic profiles of non-conventional yeasts in liquid culture.**

A

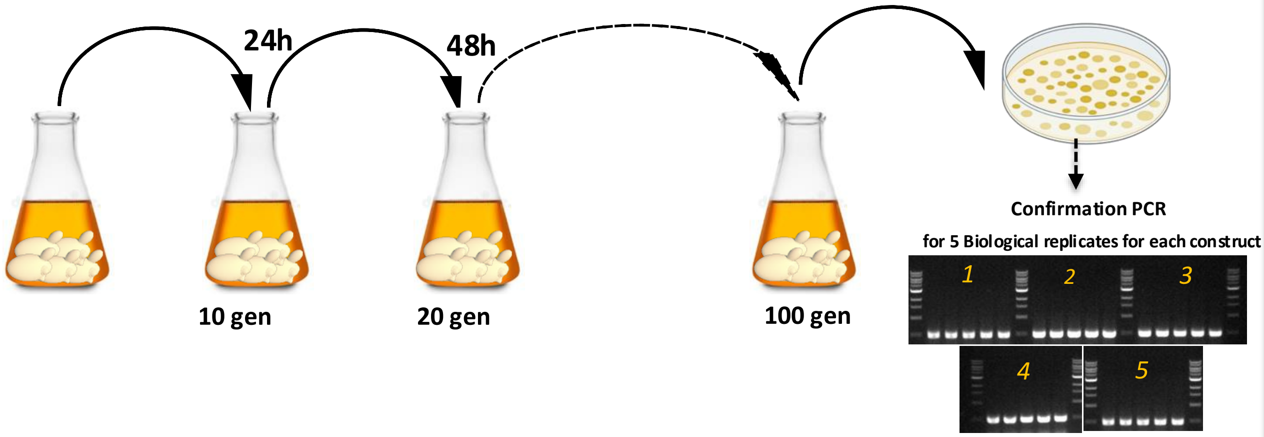


1. *S.bombicola–ve control*
2. *S.bombicola* +POT2-G418-pan/ARS-*TDH3p* (*S. c*)–*ldh.F* (*L. h*)–*TDH3t* (*S. c*)
3. *S.bombicola* +POT2-G418-pan/ARS-*TDH3p* (*M. b*)–*ldh.F* (*L. h*)–*TDH3t* (*M. b*)
4. *S.bombicola* +POT2-G418-pan/ARS-*TDH3p* (*S. c*)–*ldh.F* (*B. M*)–*TDH3t* (*S. c*)
5. *S.bombicola* +POT2-G418-pan/ARS-*TDH3p* (*M. b*)–*ldh.F* (*B. M*)–*TDH3t* (*M. b*)

**Supplementary Figure 2. Plasmid stability of the broad-host expression vector harbouring *ldh***

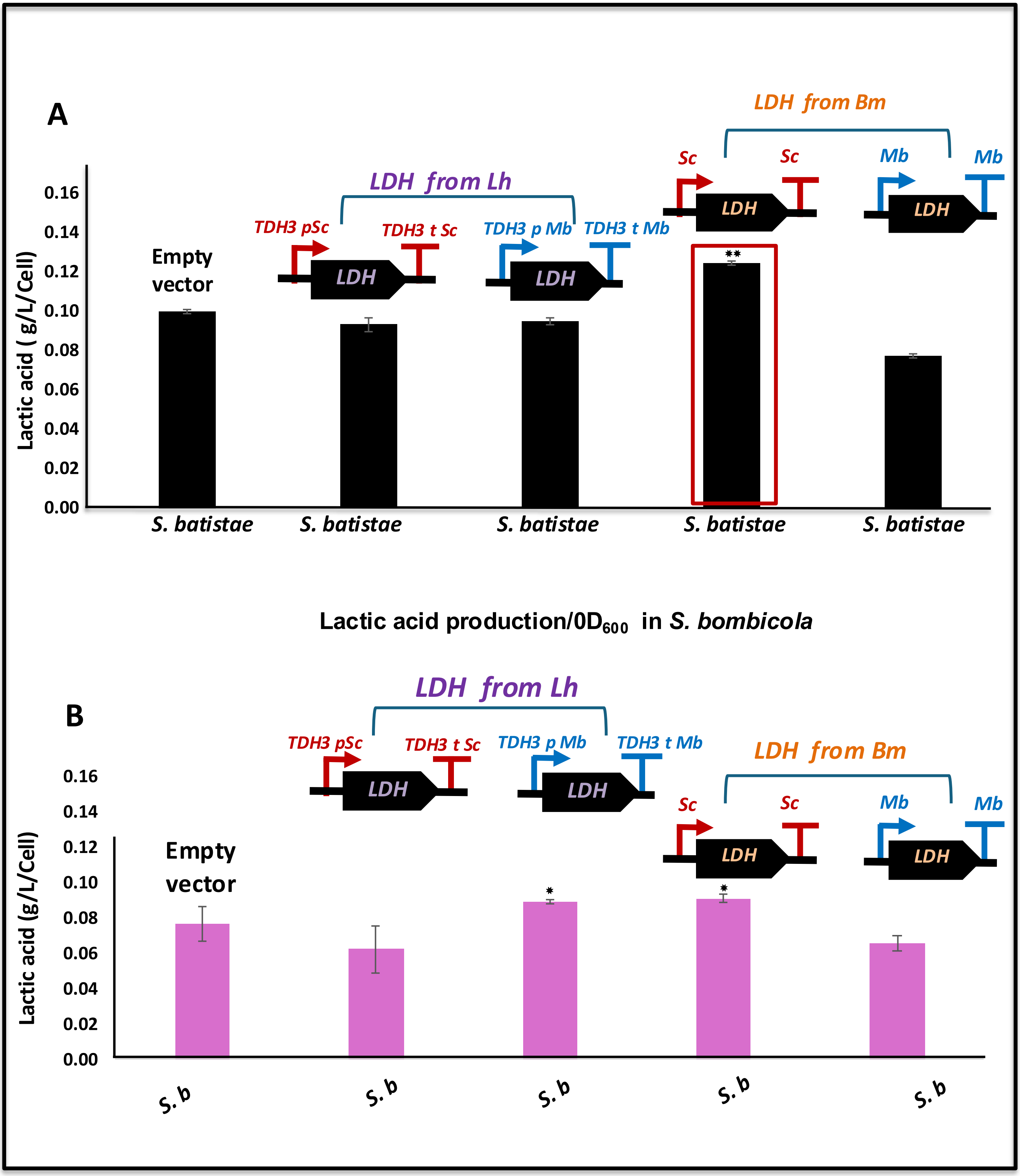


**Supplementary Figure 3. Biomass-normalised lactic acid production reveals species-specific responses to heterologous *ldh* expression in non-conventional yeasts.**

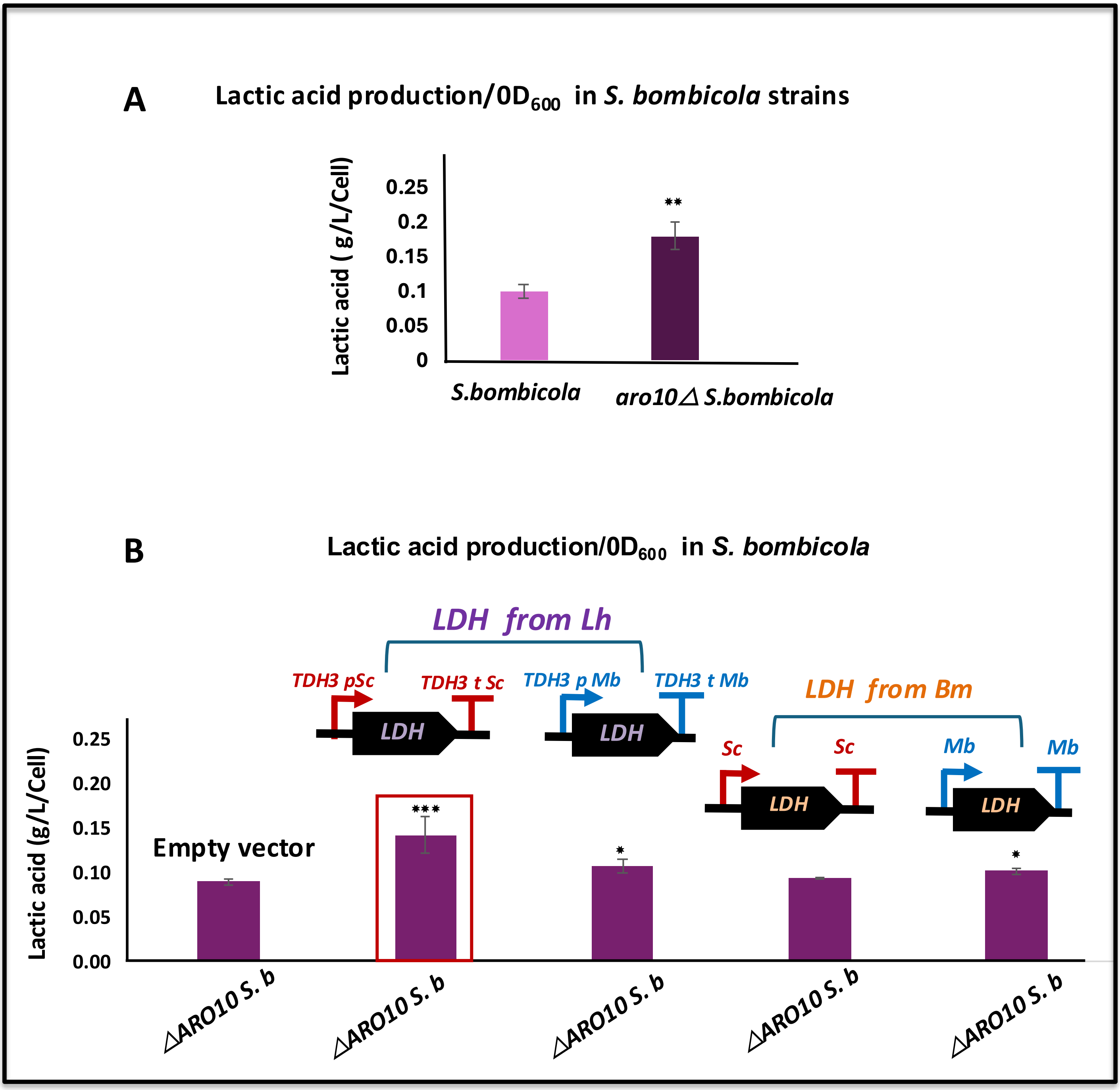


**Supplementary Figure 4. Deletion of *ARO10* redirects carbon flux and enhances endogenous and heterologous lactic acid production in *S. bombicola, normalised to $OD_{600}$***
